# Integrative genomics uncover mechanisms of renal medullary carcinoma transformation, microenvironment landscape and therapeutic vulnerabilities

**DOI:** 10.1101/2021.09.29.462391

**Authors:** Bujamin H. Vokshi, Guillaume Davidson, Alexandra Helleux, Marc Rippinger, Alexandre R. Haller, Justine Gantzer, Jonathan Thouvenin, Philippe Baltzinger, Rachida Bouarich, Valeria Manriquez, Sakina Zaidi, Pavlos Msaouel, Xiaoping Su, Hervé Lang, Thibault Tricard, Véronique Lindner, Didier Surdez, Jean-Emmanuel Kurtz, Franck Bourdeaut, Nizar M. Tannir, Irwin Davidson, Gabriel G. Malouf

**Affiliations:** Department of Cancer and Functional Genomics, Institute of Genetics and Molecular and Cellular Biology, CNRS/INSERM/UNISTRA, 67400 Illkirch, France; Department of Medical Oncology, Institut de Cancérologie Strasbourg Europe,67200 Strasbourg, France; Department of Genitourinary Medical Oncology, The University of Texas MD Anderson Cancer Center, Houston, TX 77030, USA; Department of Bioinformatics and Computational Biology, Division of Quantitative Sciences, The University of Texas MD Anderson Cancer Center, Houston, TX 77030, USA; Department of Urology, CHRU Strasbourg, Strasbourg University, 67000, Strasbourg; Department of Pathology, CHRU Strasbourg, Strasbourg University, 67200, Strasbourg; INSERM U830, Équipe Labellisée LNCC, Diversity and Plasticity of Childhood Tumors Lab, Institut Curie Research Centre, 75005 Paris, France; Balgrist University Hospital, University of Zurich, Zurich, Switzerland; INSERM, U830, Pediatric Translational Research, PSL Research University, SIREDO Oncology Center, Institut Curie, Paris, France; ‘Équipe Labellisée’ Ligue National contre le Cancer

**Keywords:** Renal medullary carcinoma, SMARCB1, TFCP2L1, MYC, ferroptosis, EMT

## Abstract

Renal medullary carcinoma (RMC) is an aggressive desmoplastic tumour driven by bi-allelic loss of SMARCB1, however the cell-of-origin, the oncogenic mechanism and the features of its microenvironment remain poorly understood. Using single-cell and multi-region sequencing of human RMC, we defined transformation of thick ascending limb (TAL) cells into at least three RMC cell states along an epithelial-mesenchymal gradient through a transcriptional switch involving loss of renal transcription factor TFCP2L1 and gain of a NFE2L2-associated ferroptosis resistance program. SMARCB1 re-expression in cultured RMC cells reactivates TFCP2L1 that relocates SWI/SNF from the promoters of the MYC-driven oncogenic program to the enhancers of TAL identity genes followed by ferroptotic cell death. We further show that RMC is associated with abundant M2-type macrophages and cancer-associated fibroblasts (CAFs) and we identify key regulatory cross-talks that shape this immunosuppressive microenvironment. Together our data describe the molecular events of RMC transformation and identify novel therapeutically targetable vulnerabilities.

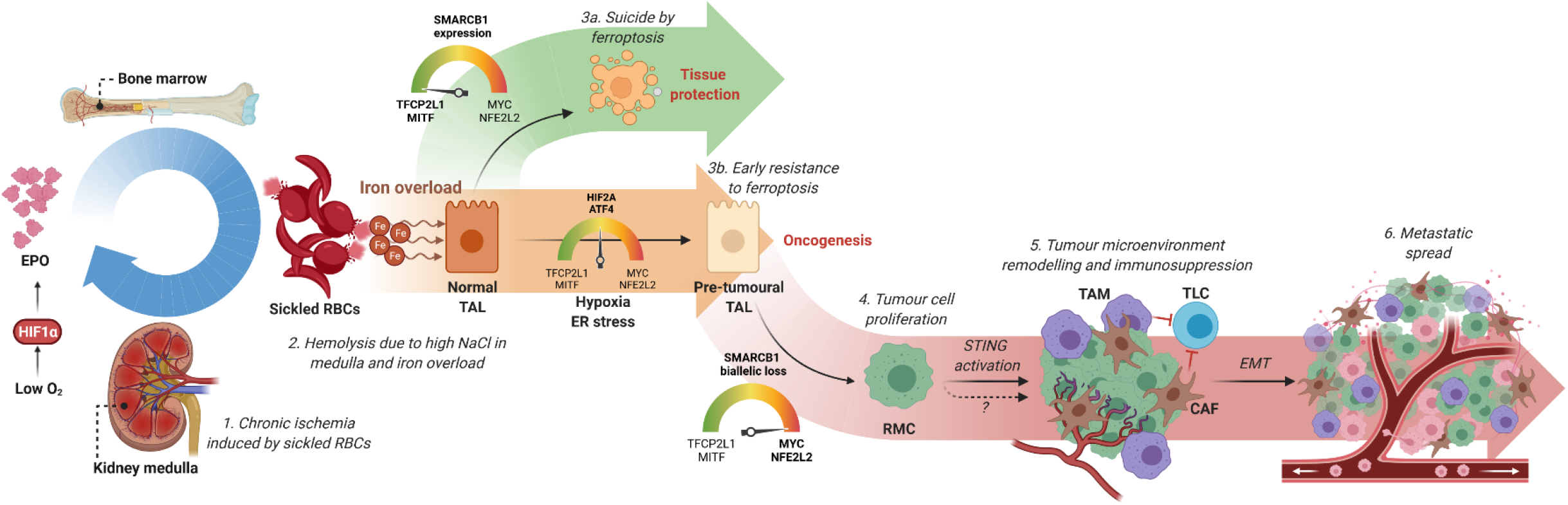

## Introduction

First described in 1995 (Davis et al., 1995), renal medullary carcinoma (RMC) is a lethal malignant neoplasm arising from the kidney medulla region. Despite its relative rarity, RMC is the third most common renal cancer among young adults (Cajaiba et al., 2018). It typically afflicts male patients of African descent with sickle cell trait at a median age of 28 years, yet the association is still poorly understood (Alvarez et al., 2015; Msaouel et al., 2019). RMC is highly aggressive with most patients presenting metastatic disease at the time of diagnosis and less than 5% survive longer than 36 months (Carugo et al., 2019; Msaouel et al., 2020a). In addition, RMC tumours are resistant to targeted therapies used for other renal cancers and the best available cytotoxic chemotherapy regimens produce a brief objective response in less than 30% of cases (Msaouel et al., 2020b; Swartz et al., 2002). Alternative treatments such as anti-angiogenics, EZH2 inhibitors and immunotherapy have been tested with varying success (Msaouel et al., 2020a). RMC tumour tissue resembles a high-grade carcinoma exhibiting reticular or cribriform patterns and usually stain positive for VIM, MUC1, pankeratins, PAX8, HIF1α and VEGF (Gupta et al., 2012; Swartz et al., 2002). RMC are also characterized by a strong desmoplasia, a prominent inflammatory infiltrate as well as the frequent presence of sickled red blood cells (Beckermann et al., 2017; Dimashkieh et al., 2003). The landscape of immune cells and cancer-associated fibroblast features in RMC tumours remains to be elucidated.

The hallmark of RMC is loss of SMARCB1 expression (Elliott and Bruner, 2019), a core subunit of the SWItch/Sucrose Non-Fermentable (SWI/SNF) chromatin remodelling complex. Several mechanisms lead to SMARCB1 loss in RMC including deletions, point mutations, inactivating translocations and loss-of-heterozygosity (Msaouel et al., 2020a). SMARCB1 loss is also the hallmark of malignant rhabdoid tumours (RTs), atypical teratoid/rhabdoid tumours (ATRTs) and epithelioid sarcomas (ESs). The majority of RTs and RMCs share common features such as their renal location and low mutation burden (Msaouel et al., 2020a). We recently characterized the molecular characteristics of RMC identifying frequent chromosome 8q gain associated with a copy-number gain of MYC (Msaouel et al., 2020a). SMARCB1 loss activates the MYC pathway resulting in increased DNA replication stress and DNA damage response. RMC are thought to arise from the distal region of the nephron, however evidence is limited to correlation inference using bulk RNA-seq data from 8 nephron biopsies with identified renal cell populations (Cheval et al., 2012; Msaouel et al., 2020a). Thus, despite the above pathology and molecular characterization, the cell of RMC origin is as yet not fully defined and the molecular mechanisms involved in oncogenic transformation associated with SMARCB1 loss remain poorly characterized.

To address these issues, we integrated data from single-cell (sc)RNA sequencing of human tumours, multi-region RNA sequencing, bulk transcriptomic data from 2 RMC cohorts, and SMARCB1 gain of function experiments in cellular models. This comprehensive approach revealed how the thick ascending limb (TAL) cells are transformed into RMC through a transcriptional switch involving loss of renal master regulator TFCP2L1 and activation of a NFE2L2 and GPX4-associated ferroptosis resistance program. We further defined the RMC microenvironment as immunosuppressive and characterized by high infiltration of cancer-associated fibroblasts (CAFs) and tumour-associated macrophages (TAMs), identifying novel therapeutically targetable tumour and TME vulnerabilities for this aggressive tumour.

## Results

### RMC ontogeny and molecular characterization of tumour cell states

To characterize the molecular features and ontogeny of RMC, we performed scRNA-seq on a post-treatment primary nephrectomy from an RMC patient with lung metastases at diagnosis. The patient showed complete response following 6 cycles of Methotrexate, Vinblastine, Doxorubicin, Cisplatin (MVAC) treatment. A total of 996 cells from the residual tumour site and 1722 cells from normal adjacent renal tissue (NAT) were aggregated and analysed. Seurat UMAP clustering revealed 14 distinct populations amongst which were 7 renal epithelial clusters and 7 stromal and immune clusters (Fig. 1A-B). Epithelial clusters comprised 6 groups of cells from the proximal and distal tubules and 1 group of collecting duct cells each expressing specific markers (Fig. 1C). Amongst these, we identified thick ascending limb (TAL) cells with expression of *SLC12A1*, *EPCAM*, *CDH1* and keratin 7 (*KRT7*), consistent with previous renal scRNA-seq datasets (Lake et al., 2019; Muto et al., 2021; Young et al., 2018).

**Figure 1.**
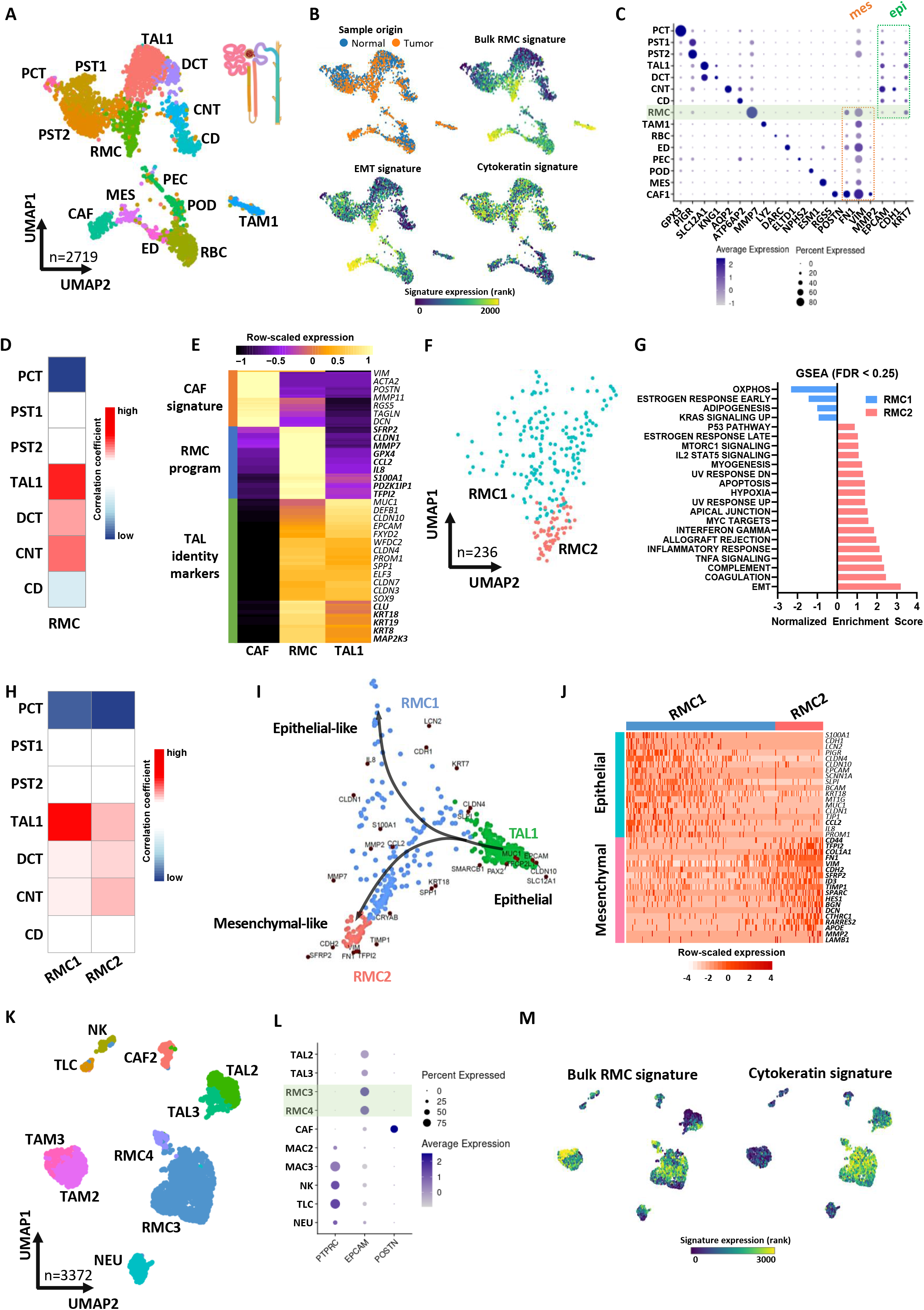
Single-cell RNA sequencing of treated (A-J) and (K-M) naive RMC tumours. **A.** UMAP plot of the aggregated treated tumour and normal adjacent tissue (NAT) representing the clusters identified by Seurat using a resolution of 1.12. *Abbreviations:* PCT: proximal convoluted tubule cells; PST1/2: proximal straight tubule cells 1 and 2; RMC: Renal medullary carcinoma cells; TAL1: thick ascending tubule cells of Henle’s loop; DCT: distal convoluted tubule cells; CNT: connecting tubule cells; CD: collecting duct cells; CAF: cancer-associated fibroblasts; MES: mesangial cells; ED: endothelial cells; RBC: red blood cells; PEC: parietal epithelial cells; POD: podocytes; TAM1: tumour-associated macrophages. **B.** UMAP projection of sample origin or selected gene signatures. **C.** Dot-plots representing gene markers of each identified cluster in the RMC treated sample. Rectangles regroup clusters according to either mesenchymal or epithelial markers. **D.** Clustifyr correlation between RMC cells and renal epithelial tubules transcriptomes. **E.** Pseudo-bulk heatmap of 100 RMC-specific and 50 CAF-specific genes using CAF1, RMC and TAL1 clusters as a matrix. **F.** UMAP representing RMC subclusters as identified by Seurat using a resolution of 1. **G.** GSEA showing enriched “Hallmark gene sets” in RMC1 relative to RMC2 cell clusters. **H.** Clustifyr correlation between RMC subclusters and renal epithelial tubules transcriptomes**. I.** SWNE trajectory analysis of the treated RMC clusters using a set of selected markers per cluster and assuming TAL1 cells as origin. **J.** Heatmap representation of a set of selected EMT genes in the 2 RMC subclusters. **K**. UMAP plot of the naive tumour cell clusters as identified by Seurat. RMC3/4: Renal medullary carcinoma cells; TAL2/3 : thick ascending tubule cells of Henle’s loop; NEU : neutrophils; CAF2 : cancer-associated fibroblasts; NK : natural killers; TLC : T-lymphocyte cells; TAM2/3 : tumour-associated macrophages. **L.** Dot-plots of selected gene markers of immune, epithelial and CAF cells. **M.** UMAP projection of the bulk RMC and cytokeratin signatures.

After merging cancer and NAT samples, we identified populations enriched in the tumour sample comprised TAMs and 2 clusters of cells harbouring an epithelial mesenchymal transition (EMT) signature which we identified to be the RMC tumour and CAF cells (Fig. 1B). All three clusters expressed specific markers (*LYZ*, *MMP7* and *POSTN* respectively) with cytokeratin expression in RMC cells (Fig. 1C). Further analyses of RMC and CAFs showed that each expressed overlapping as well as distinct sets of EMT markers (Fig. S1A and Dataset S1A).

The UMAP plot revealed that RMC cells were located close to the TAL population, consistent with a putative cell of origin located in the distal part of the nephron. We interrogated all renal epithelial populations for shared transcriptional signatures with RMC cells and found the best correlation with TAL cells of the kidney medulla (Fig. 1D). Differential gene expression analysis of a pseudo-bulk reconstitution of the RMC versus the CAF populations identified about 150 signature genes for RMC and 50 genes for CAF (Fig. 1E). RMC cells showed a specific oncogenic program, but retained many genes associated with TAL and more broadly epithelial identities (Fig. 1E). RMC and CAF cells did however commonly express EMT genes such as *VIM* and *FN1*, in contrast to TAL cells (Fig. 1C, 1E). Altogether, these observations identified TAL cells to be the normal renal population most related to RMC and hence the likely cell-of-origin.

To investigate intra-tumoural heterogeneity, we re-clustered only the RMC cells identifying 2 distinct subpopulations, RMC1 and RMC2 (Fig. 1F). Gene Set Enrichment Analysis (GSEA) revealed that RMC1 were enriched in oxidative phosphorylation (OXPHOS), whereas RMC2 were enriched in EMT, interferon gamma, inflammatory response and hypoxia (Fig. 1G). Correlation of the RMC1 and RMC2 specific signatures to those of normal tubules revealed that RMC1 partly retained a TAL signature that was reduced in RMC2 (Fig. 1H). These observations were independently confirmed by SWNE trajectory analysis that traced transformation of TAL cells to RMC1 or RMC2 (Fig. 1I). RMC cells separated into a ‘stressed’ epithelial-like phenotype with higher levels of cytokines (*IL8*, *LCN2*), keratins and epithelial markers such as *CDH1*, *CLDN1* and into RMC2 cells with higher expression of mesenchymal markers such as *SFRP2*, *CDH2* and *FN1*. Thus, this RMC tumour comprised epithelial-like RMC1 cells and mesenchymal-like RMC2 cells (Fig. 1J).

We next analysed a naive RMC sample from a primary nephrectomy of 16-year-old patient with regional lymph node and adrenal gland metastases (pT4N1M1) at presentation capturing a total of 3372 cells. Following surgery, the patient showed rapid progressive disease under adjuvant MVAC regimen. The patient was also primary resistant to durvalumab-tremelimumab immunotherapy and EZH2 inhibitor Tazemetostat leading to death within one year of diagnosis. Among 3372 captured cells, a large group of RMC cells was identified along with TAMs and other *CD45*-expressing immune cells (Natural killers, neutrophils and T-cells), *POSTN*-expressing CAFs, and an unexpected population of tumour-associated TAL cells (Fig. 1K-L). Both the RMC and TAL cells, that segregated into two closely located groups on the UMAP plot, expressed *EPCAM* as well as a cytokeratin signature (Fig. 1L-M). The TAL3 population could be distinguished from TAL2 cells by the lowered expression of the *SLC12A1*, *HOXB9* and *PAX8* renal identity markers (Fig. 2A and Dataset S1B). The RMC3 and RMC4 populations were highly similar with the smaller RMC4 cluster displaying an additional G2/M phase cell cycle signature designating them as mitotic RMC3 cells (Fig. 2A). The SWNE trajectory representation of the TAL and RMC populations illustrated the progressive loss of TAL identity markers from the most differentiated TAL2 to TAL3 with some TAL3 cells closely related to the RMC group that retained an epithelial-like signature (Fig. 2B).

**Figure 2.**
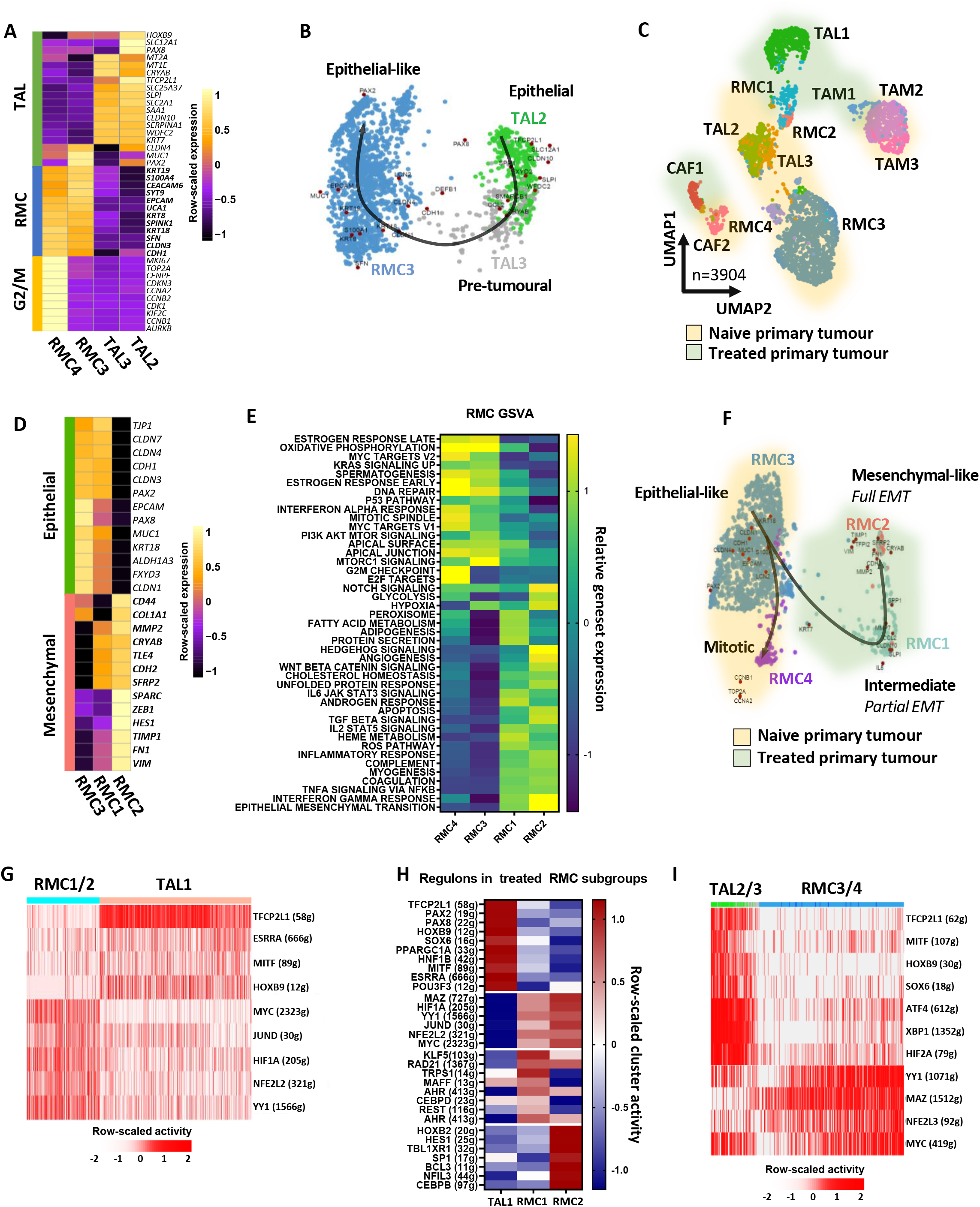
Intra-tumoural heterogeneity of RMC. **A.** Pseudo-bulk heatmap of expression of top markers of RMC and TAL clusters. **B.** SWNE trajectory analysis of the naive RMC clusters using a set of selected markers per cluster and assuming TAL2 cells as origin. **C.** UMAP representing the normalized merge of selected TAL, RMC, CAF and TAM clusters from the treated tumour (green hue) and the naive tumour (yellow hue). **D.** Pseudo-bulk heatmap showing a set of known EMT markers in all RMC clusters. Note that as RMC4 were cycling RMC3 cells, they were omitted from the analysis to avoid redundancy. **E.** GSVA analysis showing ontologies of indicated RMC clusters. **F.** SWNE trajectory analysis of normalized merged RMC clusters from treated and naive tumours using a set of differentially expressed EMT markers. **G.** SCENIC analysis of the treated tumour showing top regulons of RMC1/2 and TAL1 cells. Note that in brackets are indicated the number of genes (g) per selected regulon. **H.** SCENIC analysis of the treated tumour indicating activities of TAL regulons and RMC1- or RMC2-specific regulons. **I.** SCENIC analysis of the naive tumour revealing top regulons of RMC3/4 and TAL2/3 cells.

We then aggregated cells from the two tumours (Fig. 2C). The potential caveat of batch effect between the two samples appeared minimal as the TAM and CAF populations clustered close together. In contrast, differences between the TAL and RMC subpopulations of the 2 tumours were illustrated using a collection of epithelial, mesenchymal, endoplasmic reticulum (ER)-related stress genes (Fig. 2D, Fig. S1B). The RMC3/4 cells from the naive tumour had a marked epithelial character compared to the intermediate RMC1 cells from the treated tumour, whereas RMC2 cells had the most mesenchymal phenotype (Fig. 2D). GSVA analyses revealed enrichment for cell cycle in RMC4 cells, OXPHOS and apical junction in RMC3 cells, and EMT and interferon gamma response in RMC1 and RMC2 cells (Fig. 2E). SWNE trajectory analyses highlighted the gradient of epithelial to mesenchymal phenotypes of the different populations (Fig. 2F).

We further performed multi-region tumour RNA-seq on a cohort of four patients, for which single region transcriptome sequencing was previously reported along with that of 7 additional cases ((Msaouel et al., 2020a), designated as the MDACC cohort; and Dataset S2). Overall, we generated an additional 25 bulk RNA-seq from multiple regions of these primary tumours and the corresponding regional lymph nodes as well as 3 NATs. We then analysed intra- and inter-tumour heterogeneity using CIBERSORTx deconvolution to infer RMC1-3 composition of these sections (Fig. S1C-D). Primary tumour sections showed varying composition, some more enriched in the epithelial-like signature, others with epithelial-like and intermediate signatures, and a third group with all 3 signatures. In contrast, the lymph node metastases sections were strongly enriched in the mesenchymal-like signature. These data unravelled intra-tumour heterogeneity in RMC and the importance of tumour cells with a mesenchymal signature to metastatic progression.

We then used SCENIC regulon analyses software to identify transcriptional regulatory networks underlying the above signatures (Van de Sande et al., 2020). Comparison of the TAL and RMC populations from the treated tumour revealed a transcriptional switch from high HOXB9 and TFCP2L1 activity in TAL cells, to high MYC, HIF1A, YY1 and NFE2L2 activity in RMC cells (Fig. 2G). These data were consistent with the known MYC activity of RMC cells, whereas TFCP2L1 is a previously described determinant of the distal portion of the nephron (Werth et al., 2017). Top TAL regulons were progressively lost upon transformation into RMC1 and RMC2 populations exemplified by TFCP2L1, PPARGC1A, perhaps contributing to the OXPHOS signature (LeBleu et al., 2014), and HOXB9, whereas others like SOX9 were maintained (Fig. 2H).

Comparable observations were made between the TAL and RMC populations of the naive tumour with loss of TFCP2L1 activity and gain of MYC and NFE2L2/3 activity (Fig. 2I). Interestingly, while these TAL cells displayed TFCP2L1 activity they also showed a stress signature with prominent activity of ATF4, XBP1 and HIF2A. Moreover, they further showed YY1 and MYC activity, hallmarks of RMC cells. TAL cells from the treated tumour were derived from NAT, whereas in the naive tumour, they were tightly associated with the RMC cells in the tumour sample and showed a stressed pre-tumoural phenotype with activation of several RMC regulons. Each RMC population displayed a characteristic regulon activity such as cell cycle (BRCA1, E2F4/6) in RMC4 cells (Dimova and Dyson, 2005; Mullan et al., 2006), epithelial-like (OVOL2, ELF3) in RMC3 cells (Sengez et al., 2019; Watanabe et al., 2019) and mesenchymal-like (HES1, FOSL2) in RMC2 (Liu et al., 2015; Yin et al., 2019). Notably, activity of the PAX8 renal identity marker was strongly reduced in the RMC1 and RMC2 populations compared to RMC3 (Fig. S1E).

The role of TFCP2L1 in driving expression of epithelial genes was reinforced by analyses of the Cancer Cell Line Encyclopaedia (CCLE) showing positive correlation between TFCP2L1 (and also OVOL2) and EPCAM (Fig. S1F). Similarly, TFCP2L1 correlated with epithelia markers and anti-correlated with mesenchymal markers (Fig. S1G). In the TCGA chromophobe renal cell carcinoma dataset, originating also from distal tubules, TFCP2L1 and MITF expression correlated with that of CDH1 (Fig 1H).

The above data hence identified at least three RMC cell states, defined by distinct transcriptional signatures also found in patient tumour samples. NAT-derived TAL cells could further be distinguished from tumour-associated TAL cells that displayed a stressed, pre-tumoural phenotype in their transcriptional signatures and regulon activities.

### Molecular characterization of the RMC microenvironment

In addition to TAL and RMC cells, scRNA-seq revealed CAFs and immune infiltrate in both tumours that was dominated by TAMs. MCP counter analyses of the collection of patient sections indicated the abundant presence of fibroblasts and monocytic lineage cells in primary tumour and lymph node sections (Fig. S2B). In contrast, abundant B and T cells were found only in lymph node sections.

RMC tumours comprised 3 TAM populations. The TAM1 population displayed a pro-inflammatory M1 signature (Fig. 3A). In contrast, TAM2 and TAM3 displayed an anti-inflammatory M2 signature with high expression of known M2 markers IL10 and MAF (Conejo-Garcia and Rodriguez, 2020), that was strongest in TAM3 (Fig. 3A). Consistently with previous reports (Yunna et al., 2020), GSVA analyses revealed the strong activation of the TGF-beta pathway in the M1-type TAM1 cells and of the interferon gamma response and NF-kB pathway in the M2 type TAM3 cells, with TAM2 showing intermediate signatures (Fig. S2B). SWNE trajectory analysis further confirmed the idea that the TAM2 signature represented an intermediate state between the most polarized TAM1 and TAM3 states (Fig. 3B). This conclusion was supported by SCENIC regulon analyses that showed high activity of STAT1 and REL/NF-kB in the M2-signature TAM3 cells, whereas the TAM1 signature was associated with high activity of a several regulons associated with proliferation and EMT (E2F6, E4F1, TEAD4) in keeping with cell cycle associated terms in the GSVA analyses (Fig. 3C and Fig. S2B). The M1-type TAM1 cells were found only in the treated tumour, whereas TAM3 cells were strongly enriched in the naive tumour promoting its strongly immunosuppressive and pro-angiogenic state.

**Figure 3.**
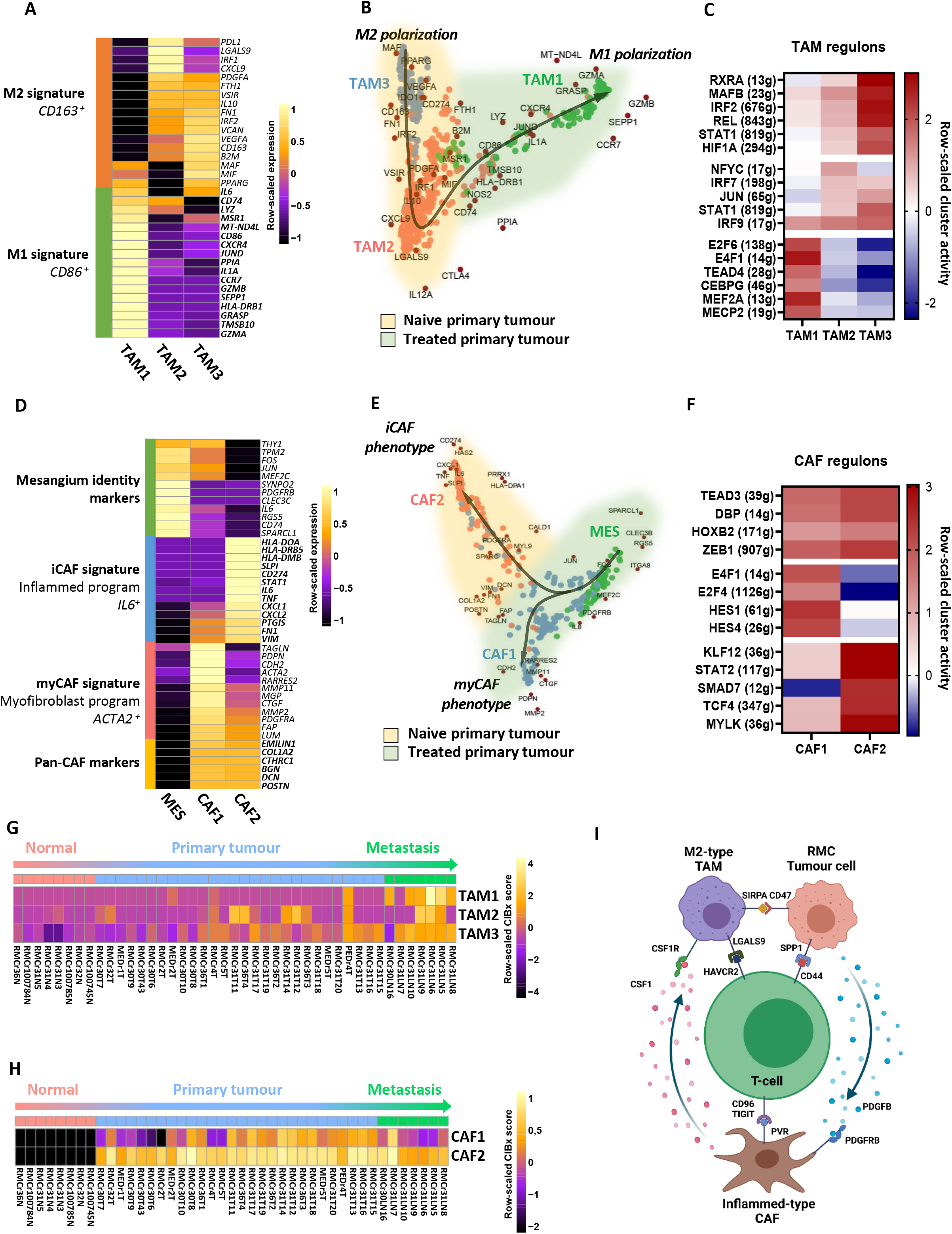
Molecular characterization of the RMC microenvironment in the treated and naive tumours. **A.** Pseudo-bulk heatmap of macrophages M1 and M2 gene signatures in TAM clusters from the treated and naive tumours. **B.** SWNE trajectory analysis of TAM clusters using a set of selected M1/M2 polarization markers revealing distinct TAM phenotypes. Arrow indicates the putative trajectory in the treated RMC sample. **C.** SCENIC analysis of normalized merged cells revealing specific regulons in each TAM cluster. **D.** Pseudo-bulk heatmap of iCAF and myCAF signature genes as well as MES and CAF identity markers in CAF clusters and their putative MES cell-of-origin. **E.** SWNE trajectory analysis using a set of selected CAF and MES markers revealing distinct CAF phenotypes. Arrow indicates the putative trajectory of CAF activation from MES cells. **F.** SCENIC analysis of normalized merged cells revealing specific regulons of each CAF cluster. **G**. Deconvolution of TAM specific signatures on bulk RNA-seq from sections of RMC primary tumours, lymph node metastasis and normal adjacent tissues. **H.** Deconvolution of CAF specific signatures on bulk RMC RNA-seq. **I.** Schematic representation of molecular cross-talks between RMC cells and their microenvironment as inferred by CellPhoneDB using our scRNA-seq.

Analogous analyses of the CAFs from both tumours revealed two populations with either a myofibroblastic myCAF signature (CAF1) predominant in the treated tumour or an inflamed iCAF signature (CAF2) found in the naive tumour (Fig. 3D). In line with previous reports (Biffi et al., 2019; Elyada et al., 2019), GSVA analyses revealed enrichment of interferon and inflammatory responses in the CAF2 population (iCAF), in contrast to protein secretion and TGF-beta signalling in CAF1 cells consistent with their myofibroblastic identity (Fig. S2D). Renal CAFs may arise from the pericyte-like mesangial cells (LeBleu and Kalluri, 2018) and SWNE analyses incorporating NAT-derived mesangial (MES) cells supported the idea they gave rise to the two CAF populations (Fig. 3E) (Hosaka et al., 2016). SCENIC analyses further highlighted the differential transcription factor activity driving the two CAF states, notably STAT2 in iCAFs and HES1 in myCAFs (Fig. 3F).

We then applied the identified TAM and CAF signatures to the bulk-RNA-seq data from the patient tumour sections as described above. The TAM2 and TAM3 signatures were detected in a subset of primary and metastases sections, whereas the TAM1 signature was poorly represented in the majority of primary tumour sections, but was highly enriched in the lymph node metastases sections (Fig 3G). Likewise, CAF2 cells were detected in all primary and metastases sections, whereas CAF1 were not present in all primary sections and lowly represented in metastases sections (Fig 3H).

These analyses showed that the naive tumour and untreated primary patient sections displayed a pro-tumoural, immunosuppressive microenvironment with predominantly iCAFs and M2-type TAMs. However, the MVAC-treated microenvironment was characterized by M1-type TAMs and myCAFs.

To identify cross-talks between the cell populations within the RMC microenvironment, we used CellPhoneDB to infer putative regulatory pathways via known ligand-receptor pair interactions in both naive and treated tumours (Fig. 3I) (Efremova et al., 2020). The CAF populations strongly contributed to the interactions with the other cell-types (Fig. S3A). Amongst the many potential interactions, CAF-derived CXCL12 could signal to its CXCR4 receptor on TAMs acting as a chemoattractant and potential mechanism for high TAM infiltration (Fig. S3B-C) (Li et al., 2019). CAFs also expressed CSF1 that signals through CSF1R to differentiate monocytes into M2-polarized TAMs (Cannarile et al., 2017). CSF1R further showed strong correlation in MDACC cohort RNA-seq data with its ligand CSF1 as well as other markers of immune suppression (Fig. S3D). We also identified CD47-expressing RMC3 cells that can interact with SIRPA on TAMs eliciting an inhibitory signal for cell phagocytosis as shown by others (Braun et al., 2021; Morrissey et al., 2020). RMC cells may signal to T-cells via SPP1-CD44 interaction described to inhibit T-cell activation (Klement et al., 2018). Expression analysis of known immune checkpoints revealed only low levels of classically targeted CTLA4 and PD1 molecules (Fig. S3E). Instead, RMC tumours displayed expression of alternative co-inhibitory signals such as TIGIT or CD96, two receptors of PVR (CD155) expressed by inflamed-type CAFs (Fig. 3I) (Gorvel and Olive, 2020).

Together these data identified a collection of potential signaling pathways by which RMC, TAM and CAF cell populations shape a strong immunosuppressive microenvironment with expression of several known immune checkpoints contributing to immune escape.

### Cell state and microenvironment of a patient derived RMC xenograft

We further analysed a patient derived xenograft (IC-PDX-132) from an RMC tumour treated with 6 cycles of cisplatin, gemcitabine and bevacizumab that had undergone 4 passages of subcutaneous injections on immunocompromised mice. Around 10,000 cells were captured and the sequences aligned to a human-mouse hybrid genome. A large group of human RMC tumour cells were identified with high expression of EPCAM and the bulk RMC signature as well as a group of murine cells corresponding to CAFs and pericytes, TAMs and monocytes, and a smaller number of other immune cells (Fig. S4A-C and Dataset S1C). A third group that we tagged ‘LQ’ (low-quality) comprised cells with high levels of mitochondrial genes and potential doublets, that were removed from the subsequent analyses. SWNE analysis traced the trajectory of monocytes into 2 TAM populations (TAM4 and 5) with murine M1 or M2-type polarization signatures, respectively (Fig. S4D). SCENIC analyses of these populations revealed transcription regulatory programs reminiscent of the human TAMs with activity of E2F factors in the M1 state and JUN and REL in the M2 state (Fig. S4E). Further SWNE analyses traced the trajectory of pericytes into myCAF-type CAF4 cells through an intermediate CAF3 population, whereas no iCAF-type cells were detected (Fig. S4F). SCENIC identified regulons active in pericytes and CAFs as well as those specific to each cell types such as TWIST1 in myCAFs (Fig S4G). Moreover, the expression of several ligand-receptor pairs seen in human RMC such as CSF1-CSF1R, CXCL12-CXCR4 were detected in the PDX (Fig. S4H).

Re-clustering the RMC cells revealed 4 subpopulations together with some mouse cells of undefined identity that were not further considered (Fig. S4I). The RMC8 cluster showed a strong cell cycle signature and regulon activity designating them as mitotic RMC cells, whereas RMC6 cells displayed high hypoxia and stress-associated regulons such as ATF4 and DDIT3 (Fig S4I-K) (Senft and Ronai, 2015). RMC5 and RMC7 on the other hand corresponded to epithelial-like and intermediate state cells respectively analogous to the RMC3 and RMC1 cells in the human tumours (Fig. S4J). No distinct highly mesenchymal population was observed, although the mitotic RMC8 cells showed the most dedifferentiated phenotype and highest expression of FN1 and CD44. SCENIC analyses of these populations identified the key MYC, YY1, and NFE2L2 regulons in the RMC cells as seen above in primary human tumours (Fig S4K).

These analyses revealed that the RMC PDX comprised principally epithelial-like, intermediate and mitotic RMC cells as well as a subpopulation of hypoxic cells consistent with the idea that angiogenesis could not fully irrigate the rapidly proliferating tumour. More strikingly however, the RMC PDX cells recreated a murine-derived microenvironment dominated by CAFs and TAMs analogous to human RMC with conservation of key molecular cross-talks.

### Cultured RMC cells recapitulate the epithelial and mesenchymal states

To better define the mechanism by which SMARCB1 loss acts to drive the transition from the TFCP2L1-TAL identity program to a MYC-driven oncogenic program seen above, we used 2 independently generated RMC cell lines, RMC2C and RMC219 (Dong et al., 2017; Msaouel et al., 2020a). RMC219 cells displayed a regular rounded morphology similar to primary RPTEC renal epithelial cells (Fig. 4A). RMC2C cells were larger with a more mesenchymal morphology and were much more invasive than the RMC219 cells (Fig. 4A, 4B). Similarly, flow cytometry indicated that RMC219 cells were EPCAM high, whereas RMC2C cells were CD44 high (Fig. 4C), a marker of RCC aggressiveness (Zanjani et al., 2018). RNA-seq identified an extensive set of differentially expressed genes confirming preferential expression of epithelial markers in RMC219 cells and mesenchymal markers in RMC2C cells (Fig 4D). GSEA revealed enrichment of EMT, inflammatory response and hypoxia in RMC2C cells, as seen in the RMC2 tumour population, and enrichment of cell cycle and DNA repair in RMC219 cells (Fig. 4E). Moreover, while OXPHOS was enriched in RMC219, glycolysis was enriched in RMC2C suggesting a metabolic switch upon EMT. MITF and POU3F3, previously reported determinants of nephron morphogenesis and TAL cell differentiation (Nakai et al., 2003; Phelep et al., 2017), were preferentially expressed in RMC219 cells, whereas EMT-transcription factors like TWIST1/2 and SNAI2 were preferentially expressed in RMC2C cells (Fig. 4F). Immunoblot analyses confirmed higher expression of VIM, and SNAI2 in RMC2C and higher expression of CDH1 and MITF in RMC219 cells (Fig. 4G). Both cell lines however showed expression of NFE2L2 and MYC and lacked SMARCB1. These cell lines therefore reproduced epithelial-like and mesenchymal-like phenotypes analogous to those observed in human tumours.

**Figure 4.**
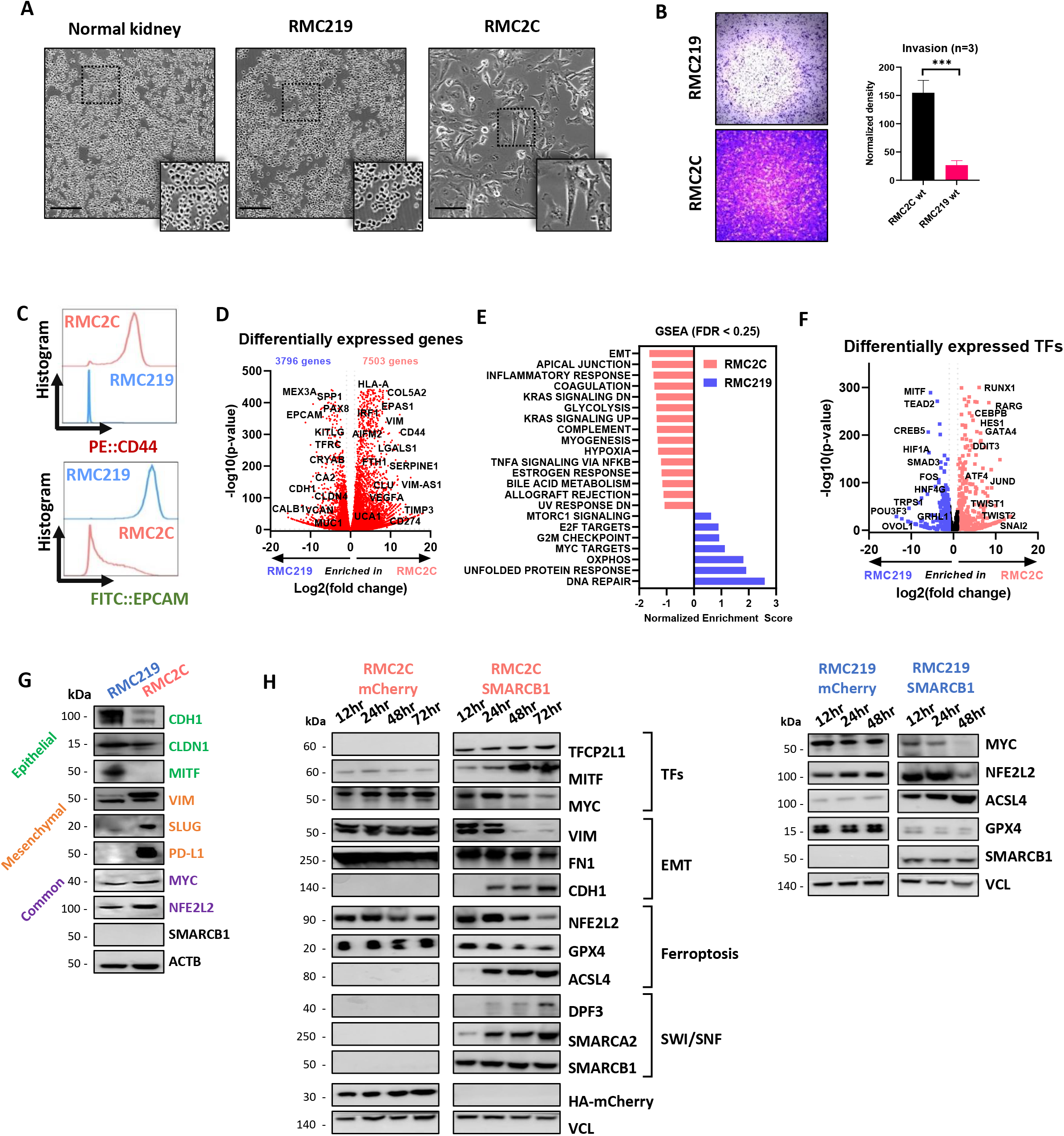
Cultured RMC cells recapitulate EMT cell states. **A.** Phase-contrast microscopy at 20X magnification of normal kidney (RPTEC) and tumour cells (RMC219 and RMC2C) showing distinct morphologies of RMC lines. **B.** Brightfield microscopy at 4X magnification of Boyden chamber matrigel assays using RMC lines (left) and absolute quantification using absorbance of resuspended crystal violet (right). **C.** Flow cytometry of membrane protein expression of EPCAM and CD44 in RMC lines. **D.** Volcano plot depicting differentially expressed genes using normalized bulk RNA-seq of RMC lines. **E.** GSEA using the Hallmarks genesets showing pathways enrichment in respective RMC lines. Note that only pathways with FDR < 0.25 are shown. **F.** Volcano plot of differentially expressed 1681 FANTOM5-defined TFs using normalized bulk RNA-seq of RMC lines. **G.** Immunoblots detecting the indicated proteins. **H.** Immunoblots showing expression of selected proteins upon re-expression of SMARCB1 in RMC2C (left) and RMC219 (right).

### SMARCB1 re-expression in RMC cells represses the oncogenic program

We analysed expression of SWI/SNF subunits in RMC2C cells compared to other SMARCB1-deficient cell lines and HEK293T kidney cells. As expected SMARCB1 was absent from all lines (Fig. S5A). The catalytic ATPase subunit SMARCA2 (BRM) was absent in all lines except VA-ES-BJ (epithelioid sarcoma), while SMARCA4 (BRG1) was detected in all lines except CHLA-06-ATRT (rhabdoid tumour). RMC2C cells showed the most important changes in SWI/SNF composition with absence of SMARCD3, ARID2 and lowest expression of DPF3, PBRM1, BRD7 and ARID1A. Although the bulk patient RNA-seq data also comprised signal from CAF and TAM cells, RMC-specific reductions in SMARCA2, and DPF3 expression could still be observed (Fig. S5B).

We engineered RMC2C and RMC219 cells to re-express SMARCB1, or mCherry as control, in a doxycycline (Dox)-dependent manner. SMARCB1 was maximally expressed in both RMC cell lines already 12 hours after Dox addition (Fig. 4H). The renewed presence of SMARCB1 induced rapid re-expression of SMARCA2, but slower re-expression of DPF3. Similarly, the TAL-associated TFCP2L1, MITF and CDH1 were also induced, whereas MYC, NFE2L2 and EMT markers VIM and FN1 were down-regulated. SMARCB1 re-expression therefore reversed key transcriptional changes observed during TAL to RMC transformation where TFCP2L1 and MITF were lost while MYC and NFE2L2 were activated.

To globally assess gene expression upon SMARCB1 re-expression, we performed RNA-seq in each cell line 12 and 48 hours after Dox-treatment. In RMC2C cells, a rapid transcriptional response was seen with 1364 up-regulated and 938 down-regulated genes after 12 hours compared to RMC219 cells where only 12 genes were up-regulated over the same period (Fig. 5A, Dataset S3). After 48 hours, a larger number of up and down-regulated genes were observed in both cell lines (Fig. 5B). Despite the differences in kinetics and numbers of affected genes, GSEA analyses revealed that in both lines, genes down-regulated were involved in oncogenic functions such as cell cycle and proliferation, designated by the GSEA terms MYC or E2F-targets, whereas up-regulated genes were designated by epithelial-like program terms such as cell adhesion, apical junction and apical surface (Fig. 5C). Comparison with bulk-RNA-seq from RMC patients relative to their NAT from both MDACC and Institut Curie cohorts showed the opposite profile with genes up-regulated in the SMARCB1-deficient tumours enriched in proliferation, cell cycle and JAK-STAT3 pathway, whereas those down-regulated associated with apical surface (Fig. 5D). Similarly, while OXPHOS was increased upon SMARCB1 expression in cell lines, it was reduced in RMC tumours. RMC cell lines hence reproduce phenotypes and transcriptional signatures seen in RMC tumours whose key features were reversed by SMARCB1 re-expression.

**Figure 5.**
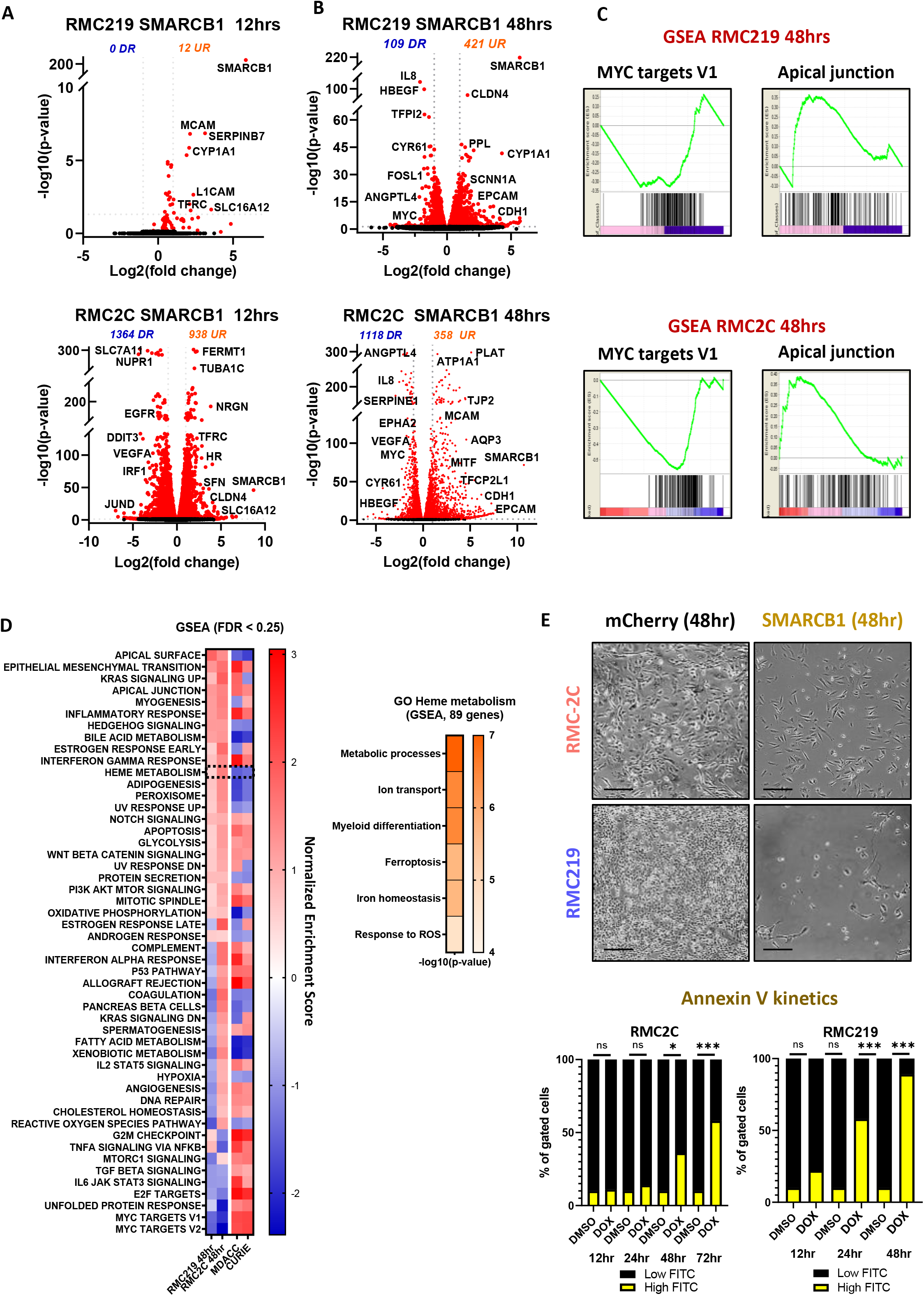
Tumour-suppressor function of SMARCB1. **A.** Volcano plot revealing up- and down-regulated genes at 12 hrs after SMARCB1 re-expression in RMC lines. **B.** Volcano plot revealing up- and down-regulated genes at 48hrs after SMARCB1 re-expression in RMC lines. **C.** GSEA showing top up- and down-regulated pathways upon SMARCB1 re-expression (48 hrs) with similar ontologies observed in both lines. **D.** Integrative heatmap showing GSEA Hallmarks enrichments (left panel) in SMARCB1 re-expressing RMC lines and 2 cohorts of RMC primary tumours (MDACC: n=11; Curie: n=5) and Metascape ontology analysis of genes constituting the GSEA “Heme metabolism” term (right panel). **E.** Phase-contrast microscopy at 10X magnification of RMC lines 48 hrs after re-expression of either SMARCB1 or mCHERRY control. Quantification of cell death in RMC lines at selected time-points upon SMARCB1 re-expression, as assessed by flow cytometry (lower panel). Note that the % of cells staining positive for either ANXA5 or propidium iodide were tagged as ‘dead’. The remaining unstained cells were tagged ‘viable’. Represented values are the mean of three biological replicates and unpaired t-test analyses were performed by Prism 5 by comparing matched time-points. P-values: ns= p>0,05; *= p<0,05; **= p<0,01; ***= p<0,001 and ****=p<0,0001.

### SMARCB1 re-expression in RMC cells induces ferroptotic cell death

SMARCB1 re-expression induced cell death with a 10-20-fold increase in the number of Annexin V-expressing cells (Fig. 5E). RMC219 cells responded rapidly with many dead cells detected by 24 hours after Dox addition, whereas death of RMC2C cells took 48 hours. To better understand the mechanism underlying cell death, we examined in more detail the gene expression changes and noted that Heme metabolism was amongst the pathways strongly up-regulated upon SMARCB1 re-expression and down-regulated in RMC patients (Fig 5D). The Heme metabolism GSEA term covers iron homeostasis, response to reactive oxygen species and ferroptosis (Fig. 5D, right panel). Following SMARCB1 re-expression, key anti-ferroptosis genes such as NFE2L2, NUPR1 and their target GPX4, a well-characterized inhibitor of lipid peroxidation (Dixon et al., 2012) were down-regulated in both lines (Fig. 6A and Fig 4H). On the other hand, Transferrin (*TF*) and transferrin receptor (*TFRC*) regulating iron uptake were both rapidly induced in RMC219 and RMC2C cells (Fig. 5A-B, and 6A). Following these acute events, at 48 hours we observed increased expression of a subset of genes involved in lipid peroxidation namely DPP4, LOX, LPCAT paralogs and ACSL4 (Fig. 6A and Fig 4H). These data suggested that SMARCB1 re-expression induced an acute increase in iron uptake followed by increased lipid peroxidation and ferroptosis.

**Figure 6.**
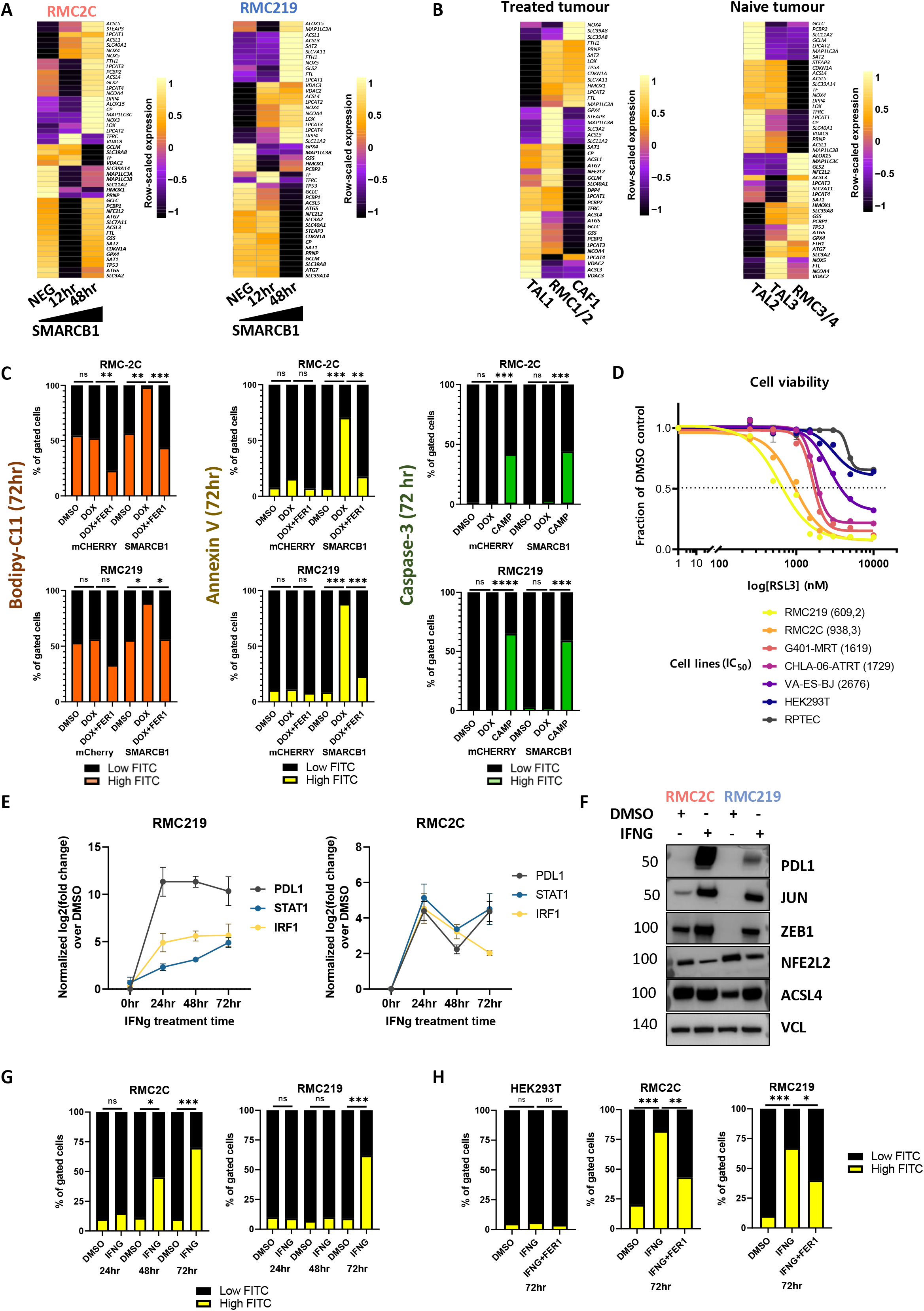
SMARCB1 regulates ferroptosis. **A.** Heatmap showing the KEGG ferroptosis gene signature in SMARCB1 re-expressing RMC2C (left) and RMC219 (right) cells. **B.** Heatmap showing expression of the ferroptosis gene signature in RMC and TAL clusters. **C.** Flow cytometry quantification of Bodipy-C11, ANXA5 and cleaved CASP3 at 72hrs in SMARCB1 or mCHERRY expressing cells and using either Ferrostatin-1 (Fer1) or camptothecin (CAMP) as controls. **D.** Cell viability (IC50) upon increasing concentrations of RSL3, a class II ferroptosis inducer (FIN). Represented values are the mean of four biological replicates of Prestoblue immunofluorescence staining. **E.** Gene expression changes of known IFNg downstream targets upon treatment of RMC lines with 10ng/mL recombinant human IFNg. **F.** Immunoblots showing expression of selected EMT and ferroptosis markers in RMC lines treated either with IFNg or DMSO vehicle control. **G.** Cell death quantified by flow cytometry using ANXA5 in RMC lines. **H.** Flow cytometry-based quantification of cell death at 72hrs upon treatment with IFNg alone, IFNg with Fer1 or DMSO in RMC lines and normal kidney cells as control. Represented values are the mean of three biological replicates and unpaired t-test analyses were performed with Prism5 by comparing conditions to matched DMSO. P-values: ns= p>0,05; *= p<0,05; **= p<0,01; ***= p<0,001 and ****=p<0,0001.

Complementary observations were made from our scRNA-seq dataset where SCENIC showed that RMC tumour cells were characterized by the activation of the NFE2L2/3 regulon a major regulator of ferroptosis (Figs. 2G, 2I) (Chen et al., 2021; Dai et al., 2020). Consequently, expression of NFE2L2, GPX4 and other anti-ferroptosis genes was upregulated in RMC cells from the MVAC-treated tumour compared to TAL cells, whereas many pro-ferroptosis genes were higher expressed in TAL cells (Fig 6B). Similarly, in the naive tumour, GPX4 and anti-ferroptosis genes were upregulated in RMC compared to TAL cells (Fig. 6B). However, in agreement with their pre-tumoural phenotype, the RMC-associated TAL3 cells showed up-regulated expression of anti-ferroptosis genes and down-regulated expression of the pro-ferroptosis genes compared to the TAL2 cells. Activation of the MYC and NFE2L2/3 regulons in these cells was therefore accompanied by activation of the ferroptosis resistance program.

We next assessed if SMARCB1 re-expression and increased expression of the lipid peroxidation genes translated into an elevation of lipid ROS assessed using BODIPY-C11-based flow cytometry (Fig. 6C). SMARCB1 re-expression induced a strong increase of lipid ROS in both lines not seen in mCherry control lines. High lipid ROS was associated with increased AnnexinV-positive cells. Importantly, the increase in lipid ROS and in Annexin-V positive cells were both impaired by ferrostatin-1, a known ferroptosis inhibitor (Fig. 6C). In contrast, SMARCB1 expression did not induce the activated Caspase 3 apoptosis marker unlike Campothecin treatment. To further confirm ferroptotic cell death, we treated RMC cells with the GPX4 inhibitor RSL3. The RMC cells had IC50 values 2-4 times lower than other RT cell lines and more than 100-fold lower than control RPTEC or HEKT cells (Fig 6D). These results confirmed that RMC cells were highly sensitive to GPX4 inhibition and that cell death was due to ferroptosis and not apoptosis.

IFNγ, secreted by the immune microenvironment in tumours *in situ*, induces tumour cell dedifferentiation and ferroptotic cell death in melanoma (Tsoi et al., 2018; Wang et al., 2019). IFNγ treatment of RMC219 and RMC2C resulted in durable expression of PDL1, STAT1 and IRF1 and of mesenchymal markers JUN and ZEB1, induced in RMC219 cells and up-regulated in RMC2C cells (Fig. 6E-F). In contrast, NFE2L2 expression was reduced. IFNγ treatment induced death of RMC2C cells between 48 and 72 hours, whereas death of RMC219 cells required 72 hours (Fig 6G). Importantly, treatment with ferrostatin 1 diminished the IFNγ-induced cell death showing that it involved ferroptosis (Fig. 6H), while as control no induced cell death was seen with HEK293T.

These results revealed that TAL cells were characterized by a ferroptosis sensitivity program that was progressively replaced in pre-tumoural TAL3 cells, in the RMC tumour populations and in RMC cell lines by a NFE2L2 and GPX4-high ferroptosis resistance program. This process was reversed by SMARCB1 re-expression that down-regulated NFE2L2 and GPX4 or by IFNγ treatment leading to cell death by ferroptosis.

### SMARCB1 re-expression promotes genomic SWI/SNF re-localization to enhancers with TFCP2L1 motifs

To investigate the consequences of SMARCB1 re-expression on SWI/SNF localization and the epigenome of RMC2C cells, we performed BRG1 and H3K27ac ChIP-seq 48 hours after Dox treatment of SMARCB1 or control mCherry expressing cells. RMC219 cells could not be used due to the rapid cell death upon SMARCB1 expression. SMARCB1 re-expression increased the overall number of H3K27ac peaks, but had little impact on their relative genomic distribution with similar fractions of sites at transcription start sites (TSS) and other genomic regions (Fig. S6A-B). However, comparison of read density at more than 46000 non-redundant H3K27ac sites revealed a gain of sites located distal to the TSS following SMARCB1 re-expression (cluster G2, Fig. S6C), whereas only a minor change was seen at the TSS. A fraction of gained peaks were extended regions reminiscent of super-enhancers (SE) known to regulate genes involved in critical aspects of lineage identity or oncogenic transformation (Hnisz et al., 2013; Whyte et al., 2013). While a large number of H3K27ac-marked SEs and their associated genes were shared between the mCherry and SMARCB1 expressing cells, 240 SE-associated genes were specific to the mCherry line and associated with a variety of functions notably DNA repair and cell cycle (Fig. S6F). More strikingly, 330 SE-associated genes specific to SMARCB1 expressing cells were associated with kidney epithelium development and differentiation as well as cell polarity and junction (Fig. S6G).

SMARCB1 re-expression also modified BRG1 genomic occupancy with a loss mainly at the TSS (H4, Fig. S6D), but a gain at distal sites (H8, Fig. S6D). Integration of BRG1 and H3K27ac read density profiles at more than 40,000 non-redundant co-occupied sites identified those with concomitant gain of H3K27ac and BRG1 following SMARCB1 re-expression (A3, Fig. 7A) predominantly located distal to the TSS (C2, Fig. 7A). In contrast, cluster A2 defined sites with reduced BRG1 predominantly located at the TSS (A2/B1, Fig. 7A) with a smaller set at distal sites (C1, Fig. 7A). Correlation with RNA-seq data indicated that genes associated with cluster A3/C2 showed increased expression following SMARCB1 re-expression (Fig. 7A). RSAT analyses revealed a strong enrichment for TFCP2L1, HOXB9, and MITF binding motifs at the distal gained A2/C3 sites (Fig. 7B). Moreover, ontology analyses of the nearest genes to the A3/C3 sites showed enrichment in differentiation, cell adhesion and kidney epithelium development (Fig. 7C). Enhanced BRG1 recruitment and H3K27ac modification was exemplified by the *CDH1* and *TJP2* loci where SMARCB1 re-expression led to increased H3K27ac at several putative upstream and intronic enhancers where a strong BRG1 recruitment was also seen (Fig. S6H). An analogous profile was observed at the *MITF* locus with *de novo* recruitment of BRG1 and H3K27ac modification at a set of putative enhancer elements located between the promoters of the MITF-A and B isoforms and downstream of the MITF-B TSS (Fig. S6I). SMARCB1 re-expression therefore led not only to re-expression of TFCP2L1 and MITF, but also re-localization of BRG1 and H3K27ac modification at putative distal enhancers bound by these factors and associated with an epithelial gene expression program.

**Figure 7.**
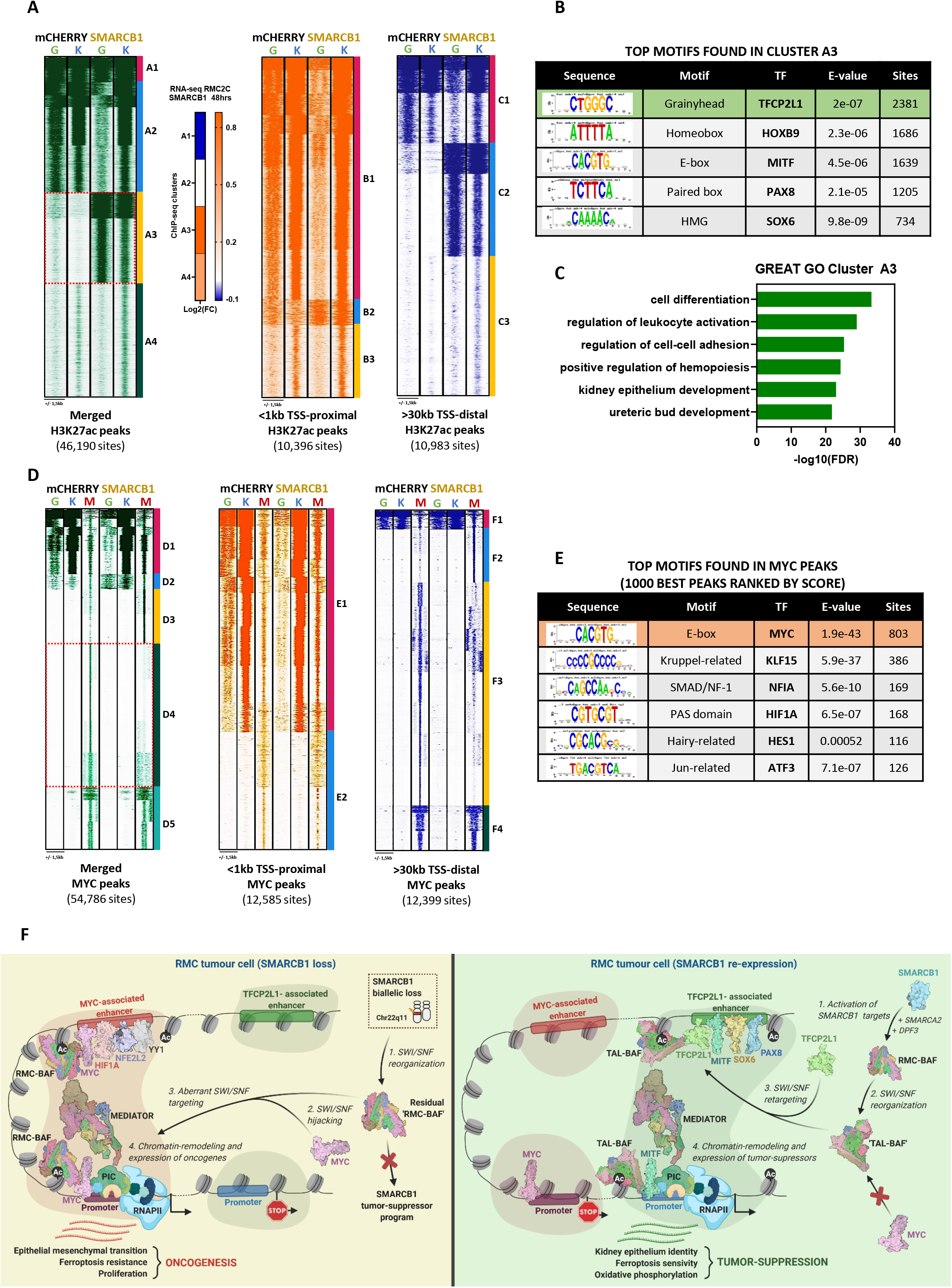
SMARCB1 retargets SWI/SNF complexes to enhancers bearing TFCP2L1 motifs. **A.** Read density maps showing genome localization BRG1 (G) and H3K27ac (K) in RMC2C cells expressing either SMARCB1 or mCHERRY using as a reference all merged H3K27ac sites (1^st^ panel), all TSS-proximal H3K27ac sites (3^rd^ panel) and all TSS-distal H3K27ac sites (4^th^ panel). Expression changes for genes associated with BRG1/H3K27ac- clusters following SMARCB1 re-expression are shown in the 2^nd^ panel. **B.** RSAT-based motif enrichment analysis using A3 sites ranked by number of sites. **C.** Ontology analysis of genes associated with A3 as annotated by GREAT. **D.** Read density maps showing genome localization of BRG1 (G), H3K27ac (K) and MYC (M) in RMC2C cells expressing either SMARCB1 or mCHERRY using as a reference all merged MYC sites (1^st^ panel), all TSS-proximal MYC sites (2^nd^ panel) and all TSS-distal MYC sites (3^rd^ panel). **E.** RSAT-based motif enrichment analysis using one thousand best MYC peaks ranked (by peak score). **F.** Working model of the oncogenic and SMARCB1 tumour-suppressor events in RMC.

### SMARCB1 re-expression remodels MYC genomic binding

As shown above, SMARCB1 re-expression diminished MYC expression. Moreover, it has been reported that SMARCB1 interacts directly with MYC to antagonize its DNA binding (Weissmiller et al., 2019; Woodley et al., 2021). To address this in a global manner, we performed MYC ChIP-seq in SMARCB1-expressing and mCherry control cells. In keeping with its reduced expression, around 50% fewer peaks were observed in SMARCB1 expressing cells compared to mCherry, but with a relative re-localization to the TSS that increased from 24% to 41% of the detected peaks (Fig. S6A-B). Read density profiles at 54,786 non-redundant MYC sites identified those with gained (I2/I8, Fig. S6E) or diminished (I3/I9, Fig. S6E) occupancy located at both TSS proximal and distal regions. Notably, integration with BRG1 and H3K27ac datasets revealed that MYC occupancy was increased at TSS proximal sites marked by H3K27ac, but characterized by diminished BRG1 occupancy (D1/E1, Fig.7D). In contrast, a large set of distal located sites were lost upon SMARCB1 re-expression (D4/F3, Fig. 7D) with a smaller number showing increased occupancy (D3/F2). Global analyses confirmed that BRG1 flanking a subset of MYC bound sites in the mCherry control cells was diminished following SMARCB1 re-expression, whereas H3K27ac was unchanged (Fig. S7A). RSAT analysis of the top 1000 MYC peaks confirmed a strong enrichment of the cognate E-box motif (Fig. 7E).

As shown above, the term ‘MYC targets’ was a prominent hallmark of genes down-regulated by SMARCB1 re-expression. We determined the % of genes in the GSEA hallmark gene sets overlapping with those associated with each MYC sub-cluster. Genes associated with D1 sites were strongly enriched in MYC targets, mitotic spindle, mTOR, E2F. DNA repair and G2M hallmark signatures (Fig. S7B). Genes associated with D4 also displayed a similar, yet lower, enrichment in many of these pathways. In agreement with this, genes associated with D1 and D4 showed global down-regulation (Fig. S7C), whereas those associated with D2 and D3 showed up-regulated expression. Thus, many genes associated with oncogenic transformation and down-regulated by SMARCB1 re-expression were associated with a gain of promoter-proximal MYC, but reduced BRG1 binding.

A similar analysis of BRG1 sub-clusters, showed genes associated with A2 were strongly enriched in the above oncogenic-associated hallmarks (Fig. S7D). In contrast, A3 sites with strongly gained BRG1 binding were enriched in genes associated with apical junction/surface and kidney morphogenesis hallmarks, consistent with re-activation of an epithelium-identity program. We used ROSE to identify MYC-H3K27ac-marked or BRG1-H3K27ac-marked SEs in control and SMARCB1-expressing cells (S7E-F). The ontology of the SE-associated genes was consistent with a switch from MYC/BRG1 driving proliferation and oncogenesis in absence of SMARCB1 to TFCP2L1/BRG1 driving epithelium identity in presence of SMARCB1.

To better understand the paradoxical observation that MYC binding increases at down-regulated oncogenic genes, we looked more closely at the large set of diminished D4 sites associated with similar ontology terms to D1. Re-clustering of D4 identified a small number (J1, Fig. S7G) of promoter-proximal sites associated with H3K27ac and a large majority of distal sites (J2, Fig. S7G). Strikingly, a large number of genes were commonly associated with both clusters (Fig. S7H). Hence many genes of the oncogenic program had both promoter-proximal and distal MYC sites showing increased and decreased occupancy, respectively.

Overall, these results showed that despite lowered expression, SMARCB1 re-expression did not repress MYC genomic occupancy, but rather remodelled its binding profile in a manner suggesting that altered enhancer-promoter communications and loss of promoter-proximal BRG1 binding underlie reduced expression of the proliferation/oncogenic program (Fig. 7F).

## Discussion

### Oncogenic events leading to RMC

Here we present a comprehensive analysis of RMC tumour cells and their microenvironment by integrating RMC cellular models, scRNA-seq from treated and naive human tumours, a novel PDX model (IC-pPDX-132) and bulk RNA-seq from multiple sections of patient cohorts. We defined at least three RMC cell states along an epithelial-mesenchymal gradient and showed that the RMC219 and RMC2C cell lines reproduced the epithelial-like and mesenchymal states respectively. A distinct population of mesenchymal-like cells was observed in the treated tumour and the mesenchymal transcriptional signature was present in primary tumours from naive patients and was predominant in the lymph nodes. Thus, de-differentiation into this mesenchymal state is not specific to drug-treated tumours, but appears to be an intrinsic feature of RMC tumours that likely contributes to their metastatic spread.

Analyses of gene expression signatures and the underlying transcription factor activity of RMC cells compared to NAT both pointed to TAL cells as RMC cell-of-origin. TAL cells were marked by strong activity of TFCP2L1 and MITF transcription factors which were associated with the epithelial expression program. MITF may act as a tumour suppressor in RMC, in contrast to melanoma where it is described as a lineage-specific oncogene driving melanoma cell proliferation (Goding and Arnheiter, 2019). Interestingly, the human germ line MITF E318K mutation predisposes to both melanoma and to renal cell carcinoma providing further evidence that MITF normally acts as tumour suppressor in this setting (Bertolotto et al., 2011). Oncogenic TAL transformation was characterized by loss of expression and activity of TFCP2L1, MITF and HOXB9, but gain of MYC and NFE2L2 that drive proliferation and ferroptosis resistance. Further evidence for this series of oncogenic events came from the fortuitous capture of tumour-associated TAL cells that displayed a pre-transformed state retaining TFCP2L1 activity, while at the same time showing MYC and YY1 activity accompanied by a hypoxia and stress signature. The first steps of transformation therefore appear to involve a hypoxia/stress driven activation of MYC prior to TFCP2L1 loss.

SMARCB1 re-expression in RMC2C cells provided experimental mechanistic support for the above model of TAL-RMC transformation. SMARCB1 expression reactivated TFCP2L1 and MITF expression and promoted BRG1 re-localization to enhancers driving expression of an epithelial expression program that were *de novo* marked by H3K27ac and enriched in TFCP2L1 and MITF binding motifs (Fig. 7F). Unfortunately, the lack of ChIP-grade TFCP2L1 and MITF antibodies does not allow us to directly confirm their presence at these enhancers. In contrast, SMARCB1 re-expression led to reduced expression of the oncogenic MYC and NFE2L2 factors. Although MYC expression was reduced by 48 hours, genomic profiling identified sites where its occupancy was either gained or reduced. Paradoxically, while MYC binding was increased at the TSS of down-regulated genes involved in proliferation and oncogenic processes, it was lost at sites distal to these genes. Although there are clear limitations in assigning distal binding sites to regulation of a given gene, a large set of genes showed increased MYC binding at the promoter and diminished binding at distal sites suggesting the importance of enhancer-promoter communication in their activation. More importantly however, BRG1 occupancy was strongly reduced at these same TSS.

Together these observations propose a model for RMC transformation where TAL cells undergoing hypoxia/stress activate MYC and NFE2L2 to drive ferroptosis resistance allowing survival under conditions favourable to SMARCB1 loss and MYC gain (Fig. 7F). SMARCB1 loss leads to inhibition of a TFCP2L1/MITF-driven TAL identity program and hence oncogenic transformation. In RMC cells, SWI/SNF complex without SMARCB1 cooperates with MYC to drive the oncogenic program, whereas upon SMARCB1 re-expression, SMARCB1-containing SWI/SNF is re-located by TFCP2L1/MITF to enhancers driving the TAL identity program. This model differs somewhat from that proposed in studies of RT cells where it was suggested that SMARCB1 can antagonize MYC DNA binding and chromatin occupancy (Weissmiller et al., 2019). In RMC cells, this antagonism translated not as a loss of MYC binding, but rather that of SMARCB1-containing SWI/SNF at MYC occupied TSS leading to reduced expression of the corresponding genes perhaps due to altered enhancer-promoter communication. In RT cells, SMARCB1 re-expression led to SWI/SNF re-localization to what are presumed to be lineage-specific enhancers (Nakayama et al., 2017; Wang et al., 2017). However, the cell of RT origin is poorly defined, and the DNA motifs at the gained enhancers were not further informative. In contrast, in RMC, SWI/SNF re-localization could be correlated with activation of the TAL identity/epithelial program, a finding further reinforced by the reactivation of a ferroptosis sensitivity signature characteristic of TAL cells. These data reveal how MYC, NFE2L2, TFCP2L1, MITF, and possibly other TFs use SWI/SNF complexes lacking or containing SMARCB1 to drive oncogenesis or the TAL identity program.

### A potential link between RMC ferroptosis and sickle cell trait

One of the key findings of our study is activation of a ferroptosis resistance pathway in RMC cells. Analyses of gene expression signatures in scRNA-seq, patient cohort RNA-seq and the RMC cell lines defined how the ferroptosis sensitivity signature in TAL cells is replaced by a ferroptosis resistance signature in RMC cells. This process is reversed in RMC cells upon SMARCB1 re-expression leading to their ferroptotic cell death unlike other RT cells where SMARCB1 re-expression leads to cell cycle arrest (Betz et al., 2002; Nakayama et al., 2017; Wang et al., 2017). Indeed, RMC cells are more sensitive to GPX4 inhibition than other RT lines. Ferroptosis is therefore a vulnerability of RMC tumours.

The above observations may help to link the RMC oncogenic process to its association with sickle cell trait. The kidney medulla is amongst the most hypoxic micro-environments in the organism (Evans et al., 2020). Due to its central role in urine concentration, the loop of Henle is characterized by increasing osmolarity and hypoxia that are highest in the TAL region. Msaouel et al. proposed a model where the high interstitial NaCl concentration induces DNA double strand breaks (DSB), whereas microcirculatory ischemia induced by red blood cell (RBC) sickling reduces this osmolarity reactivating DSB repair in a chronic hypoxic environment by NEHJ favoring translocations and deletions, particularly in fragile regions such as chromosome 22q where the *SMARCB1* locus is located (Msaouel et al., 2018).

Our observations enrich this model by showing how iron release by RBC sickling favours ferroptosis of TAL cells and their renewal to maintain the homeostasis of the epithelium (Humphreys et al., 2008; Scindia et al., 2019). On the other hand, the early acquisition of ferroptosis resistance observed in the pre-tumoural TAL cells allows them to survive under the high NaCl and hypoxic conditions that promote error-prone DSB repair. The increased extracellular iron concentration due to the fragility of the sickled RBCs thereby acts as a selective pressure for survival of ferroptosis resistant cells in an environment propitious to the mutagenic events associated with RMC development.

### Novel therapeutically targetable RMC vulnerabilities

We also present here a comprehensive characterization of the RMC microenvironment showing the prevalence of TAM and CAF populations. Strikingly, analyses of the RMC PDX (IC-pPDX-132) revealed a similar TAM- and CAF-rich microenvironment formed by murine cells. RMC cells were hence intrinsically competent to program this microenvironment not only *in situ* in the renal medulla site of origin, but also in the heterologous murine subcutaneous site. This intrinsic ability may be related to activation of pro-inflammatory cGAS-STING signalling pathway previously described in these tumours (Msaouel et al., 2020a). Further analyses showed that the microenvironment of the naive tumour and PDX comprised iCAFs and M2-type TAMs that provide an immunosuppressive environment (Buechler et al., 2021; Labani-Motlagh et al., 2020). Moreover, the common PD1 and CTLA4 immune checkpoints were lowly expressed in RMC that consequently are poorly responsive to FDA-approved immune checkpoint inhibitors (unpublished data). We rather found that RMC tumours were frequently characterized by expression of PVR together with TIGIT and/or CD96, a recently described targetable immune checkpoint (Gorvel and Olive, 2020). RMC were also characterized by CSF1-CSFR1 expression that is also targetable by small molecular inhibitors with several CSF1R-targeting therapies currently under evaluation in the clinic (Cannarile et al., 2017). Together, our analyses of RMC and its microenvironment identified several additional vulnerabilities, GPX4 inhibitors or other pro-ferroptotic drugs, novel ICIs and CSF1R inhibitors each of which may be promising new targets for therapeutic intervention.

## Methods

### Tumour Samples

The two RMC samples subjected to scRNAseq were collected from Strasbourg University Hospital and Curie Institute, according to institutional guidelines. Sample collection for further research analysis was approved ethical Committees of participating institutions and all patients provided an informed written consent for the use of material for further research. Regarding bulk RNAseq, beside the RNAseq of 11 patients recently reported (Msaouel et al., 2020a), we generated an additional dataset of multi-region RNAseq of a cohort of 4 RMC patients, including multiple sections and lymph nodes metastasis (Dataset S2).

### Human single-cell sample preparation and RNA-seq

Following the treated tumour resection, samples from the tumour and adjacent non-malignant normal adjacent tissue were each conserved at 4°C in 1mL of MACS Tissue Storage Solution (Miltenyi Biotech). Single cell suspensions were prepared using gentleMACS^TM^ dissociator and human tumour dissociation kit (Miltenyi Biotech) following manufacturer’s instructions. Samples were applied to a MACS SmartStrainer 70µm (Miltenyi Biotech) placed on a 15mL Falcon tube and 10mL DMEM were used to wash C tube and SmartStrainer. Following centrifugation at 300g and 4°C for 10min, cells were sorted using CD45 (TIL) Microbeads (Miltenyi Biotech). CD45+ and CD45- fractions were centrifuged (300g, 10min, 4°C) and dead cells were removed using Dead cell removal kit (Miltenyi Biotech). CD45- and CD45+ were mixed in 1 to 4 ratios. Cell viability and concentration were assessed before 3’-mRNA single-cell libraries were prepared using the Chromium (10x Genomics) following the manufacturer’s instructions. Libraries were sequenced 2×100bp on HiSeq4000 sequencer.

Folowing resection of the naive tumour, the sample was cut in small pieces then dissociated 30 min at 37^°^C in CO2-independent medium (Gibco) + 0,4 g/l of human albumin (Vialebex) with Liberase TL (Roche) 150 ug/ml and DNase 1 (Sigma) 150 ug/ml. Dissociated cells were then filtered with a 40 mm cell strainer, then washed and resuspended with CO2-independent medium + 0,4 g/l of human albumin. A fraction of the cell suspension was used to enrich tumor cells using Tumor isolation kit (Miltenyi Biotech, cat#130-108-339). Cells were then resuspended at 800 cells/ul in PBS + BSA 0,04%. Tissues were processed within 1 hour after tumor resection, and sorted cells were loaded in a 10x Chromium instrument within 6 hours.

### Patient-derived xenograft sample preparation

Renal medullary carcinoma (RMC) patient derived xenograft (IC-pPDX-132) was established from a resected RMC tumour treated with 6 cycles of cisplatin, gemcitabine and bevacizumab. The undissociated tumor was engrafted in the subscapular fat pad of NSG (NOD.Cg-Prkdc^scid^ IL2^rgtm1Wjl^/SzJ) mice. A PDX tumor fragment was then serially transplanted using the same procedure into Swiss Nude (Crl:NU(Ico)-Foxn1^nu^) mice until passage 4 which was used for the single cell RNA-seq experiments. Animal care and use for this study were performed in accordance with the recommendations of the European Community (2010/63/UE) for the care and use of laboratory animals. The housing conditions were specific pathogen free (SPF) for all models. Experimental procedures were specifically approved by the ethics committee of the Institut Curie CEEA-IC #118 (Authorization APAFIS#11206-2017090816044613-v2 given by National Authority) in compliance with the international guidelines. The establishment of PDX received approval by the Institut Curie institutional review board OBS170323 CPP ref 3272; n de dossier 2015-A00464-45). Written institutional informed consent was obtained from the patient.

### scRNA-seq analysis of human primary RMC tumours

After sequencing, raw reads were processed using CellRanger (v 3.1) to align on the hg19 human genome, remove unexpressed genes and quantify barcodes and UMIs. Data were then analysed in R (v4.0.2). For the treated tumour, tumour and NAT samples were aggregated with the cellranger ‘aggr’ command. The resulting aggregation was analysed with Seurat v3.2.0 following the recommended workflow. Cells were filtered for feature count ranging from 120 to 2000 and percentage of mitochondrial reads <15%. Counts were normalized with the “LogNormalize” method and data scaled to remove unwanted sources of variation (UMI count and mitochondrial reads). The number of principal components was determined from the Jackstraw plots. Clustering was performed on variable features using the 25 most significant principal components and a resolution of 1.15. For the naive tumour, the same Seurat pipeline was performed using feature counts from 200 to 6000, mitochondrial read fraction <20% and a resolution of 1.0 using the 20 most significant principal component for the clustering. Aggregate analyses of tumours 1 and 2 was performed by merging the two R objects and using the Seurat sctransform function to normalize and scale data reducing the impact of technical factors.

### scRNA-seq analysis of patient-derived RMC xenograft

For the IC-pPDX-132 sample raw reads were aligned on an hg19-mm10 hybrid genome. Cells were filtered based on feature counts ranging from 200 to 7000 and global clustering performed with a resolution of 0.3 using the 20 most significant components. Human and Mouse cells were re-clustered separately by first filtering cells with mitochondrial read fraction >20% and then using a resolution of 0.4 with 25 principal components.

### Functional analysis using scRNA-seq data

Regulome analyses of active transcription factors were performed using the SCENIC v1.1.2.2 package (Van de Sande et al., 2020). Transcription factor activities were visualized on the UMAP using AUCell or as heatmaps using the R-package ‘pheatmap’. RMC correlations with the different renal tubule clusters were computed by Clustifyr v1.0.0 (Fu et al., 2020). Trajectory analyses were plotted and visualized using Similarity Weighted Nonnegative Embedding (SWNE) (Wu et al., 2018). Ligand-receptor interactions were inferred using the CellPhoneDB python package (https://github.com/Teichlab/cellphonedb) with the “statistical_analysis” argument and default parameters (Efremova et al., 2020). Gene set variation analysis were performed using the r-package GSVA (Hänzelmann et al., 2013). For the “bulk RMC signature”, the upregulated genes from the differential analysis of the MDACC RMC cohort (11 tumours versus 6 NAT) were selected using log2FC > 2 and FDR < 0.01 (Msaouel et al., 2020a). The TAM “M1/M2” signatures were adapted from (Braun et al., 2021). The CAF “iCAF/myCAF” signatures” were adapted from (Elyada et al., 2019). For all other signatures, gene sets were retrieved from either Hallmarks MSigDB or KEGG pathways. Gene signatures were computed and visualized on UMAPs using the R package VISION (https://github.com/YosefLab/VISION).

### Cell culture, establishment of RMC lines stably expressing SMARCB1

RMC219 cells were grown in HAM-F12/D-MEM (1:1) medium supplemented with 10% foetal calf serum (FCS), Glutamine 2mM, AANE and PS. RMC2C cells were grown in MEM medium with 10% FCS, AANE, 50ng/mL EGF and PS. RMC cells infected with lentiviral constructs were grown in respective media replacing normal FCS with tetracyclin-free FCS (Dutscher) and supplemented with G418 (300ug/mL). SMARCB1 expression was induced by treatment with either DMSO or 2µM of doxycycline.

Lentiviral pInducer20 vector was obtained from Addgene and the cDNA of either SMARCB1 or mCherry was cloned into the vector by Gateway. We then used pInducer20-mCherry or - SMARCB1 containing lentiviruses to infect 1×10^6^ RMC2C or RMC219 cells. Cells were selected using 500ug/mL G418 for a week and then maintained under these conditions.

### In vitro treatments

For ferroptosis, cells were either treated with DMSO or 2uM doxycycline alone or co-treated with 2uM doxycycline and 1uM ferrostatin-1 (SelleckChem, #S7243) for the indicated times. For the Caspase-3 assays, cells were either treated with 5uM camptothecin (SelleckChem, #S1288) for 4hr, DMSO or 2uM doxycycline for the indicated times. For the IFNγ experiments, cells were either treated with DMSO or 10ng/mL of IFNγ (Peprotech, 300-02).

### Cell death, caspase-3 and lipid peroxidation analyses by flow cytometry

Cells were harvested at the indicated times and co-stained with Annexin-V-FITC and propidium iodide following manufacturer instructions (BioLegend, #640914). To assess active Caspase-3, cells were fixed and permeabilized before incubation with the FITC-conjugated caspase-3 antibody following manufacturer’s instructions for subsequent flow cytometry analysis (Abcam, #65613). To assess membrane lipid perodixation, cells were stained using Bodipy 581/591 C11 (ThermoFisher, #D3861) following manufacturer’s instructions. To assess senescence, cells were treated with 100nM bafilomycin A1 (Sigma, #19-148) for 1hr followed by 2mM C12FDG (Invitrogen, #D2893) for 2hrs before being washed and harvested for flow cytometry analyses. All assays were analysed on a LSRII Fortessa (BD Biosciences) and data were analysed using Flowjo v6.8.

### Cell viability assay by fluorescence screening

5 x 10^3^ of indicated cell types were seeded on 96-well plates in four technical replicates on day 1. The next day, cells were treated either with DMSO control or with an increasing concentration of RSL3 (SelleckChem, 8155) ranging from 0 to 10µM. At day 3, cells were washed with PBS and stained using PrestoBlue (Invitrogen, A13261) according to manufacturer instructions before fluorescence was quantified on a multi-modal spectrometer (Berthold Mithras, LB940). IC50 values were calculated using the fraction of DMSO control.

### Immunostaining quantification by flow cytometry

Wildtype RMC219 and RMC2C cells were harvested and 1 x 10^6^ cells were resuspended in buffer A (PBS 1X, EDTA 2mM, inactivated FCS 1%) and 5uL of Human TruStain FcX (Biolegend, 422301) was added for 10 min at room temperature. Following blocking, cells were stained for 1hr with 5µL of conjugated EPCAM-FITC (Biolegend, 324203) and conjugated CD44-PE (Biolegend, 103023). Following two PBS washes, cells were resuspended in buffer A before flow cytometry on a LSRII Fortessa (BD Biosciences) and analysis using Flowjo v6.8.

### Boyden Chamber Invasion assays

Before seeding, 100ul of diluted Matrigel (1:20, 356234, Corning) was added in each insert (24-well 8um inserts, Corning) and left to dry for 2hrs at 37°C before being washed twice with PBS. Subsequently, RMC cells were harvested and 2 x 10^5^ cells and seeded in the Boyden chambers in corresponding media without serum. 24hrs later, migrated cells were fixed using PFA 4% for 10 min before being stained using Crystal violet for 10 min. Excess stain was washed 3 times in PBS before images were captured on phase contrast microscope. Quantification of migrated cells was done by resuspension of staining using 100mM acetic acid for 15min before luminescence was measured on a BioTek Luminescence microplate reader (using Gen5 software).

### RNA preparation and quantitative PCR

RNA isolation was performed according to standard procedures (Macherey Nagel RNA Plus kit). RT-qPCR was carried out with SYBR Green I (Roche) and SuperScript IV Reverse Transcriptase (Invitrogen) and monitored using a LightCycler 480 (Roche). The mean of ACTB, TBP, RPL13A and GAPDH gene expressions was used to normalize the results. Primer sequences for each cDNA were designed using Primer3 Software and are available in Table 2.

**Table 1.**
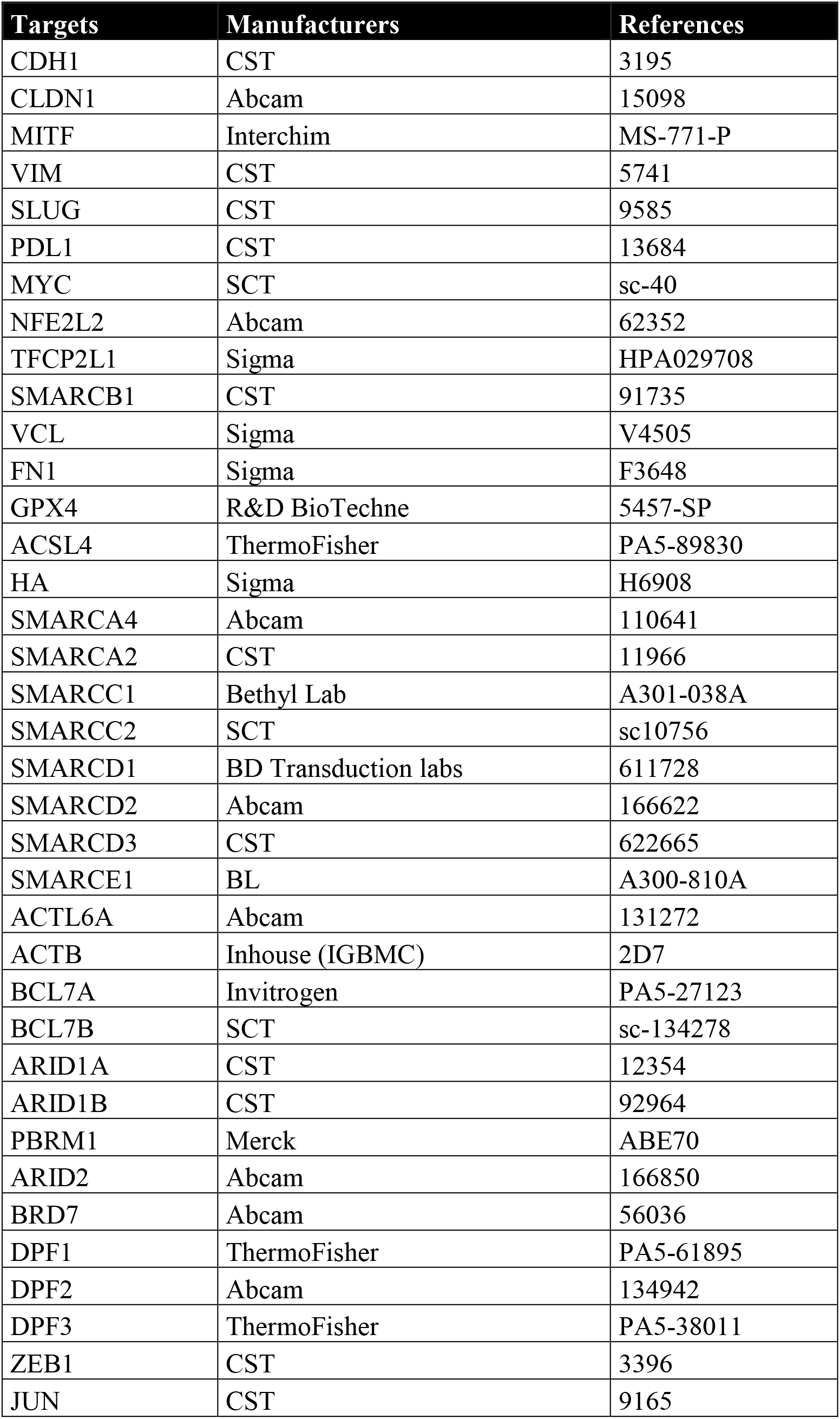
List of antibodies used.

| Targets | Manufacturers | References |
| --- | --- | --- |
| CDH1 | CST | 3195 |
| CLDN1 | Abcam | 15098 |
| MITF | Interchim | MS-771-P |
| VIM | CST | 5741 |
| SLUG | CST | 9585 |
| PDL1 | CST | 13684 |
| MYC | SCT | sc-40 |
| NFE2L2 | Abcam | 62352 |
| TFCP2L1 | Sigma | HPA029708 |
| SMARCB1 | CST | 91735 |
| VCL | Sigma | V4505 |
| FN1 | Sigma | F3648 |
| GPX4 | R&D BioTechne | 5457-SP |
| ACSL4 | ThermoFisher | PA5-89830 |
| HA | Sigma | H6908 |
| SMARCA4 | Abcam | 110641 |
| SMARCA2 | CST | 11966 |
| SMARCC1 | Bethyl Lab | A301-038A |
| SMARCC2 | SCT | sc10756 |
| SMARCD1 | BD Transduction labs | 611728 |
| SMARCD2 | Abcam | 166622 |
| SMARCD3 | CST | 622665 |
| SMARCE1 | BL | A300-810A |
| ACTL6A | Abcam | 131272 |
| ACTB | Inhouse (IGBMC) | 2D7 |
| BCL7A | Invitrogen | PA5-27123 |
| BCL7B | SCT | sc-134278 |
| ARID1A | CST | 12354 |
| ARID1B | CST | 92964 |
| PBRM1 | Merck | ABE70 |
| ARID2 | Abcam | 166850 |
| BRD7 | Abcam | 56036 |
| DPF1 | ThermoFisher | PA5-61895 |
| DPF2 | Abcam | 134942 |
| DPF3 | ThermoFisher | PA5-38011 |
| ZEB1 | CST | 3396 |
| JUN | CST | 9165 |

**Table 2.**
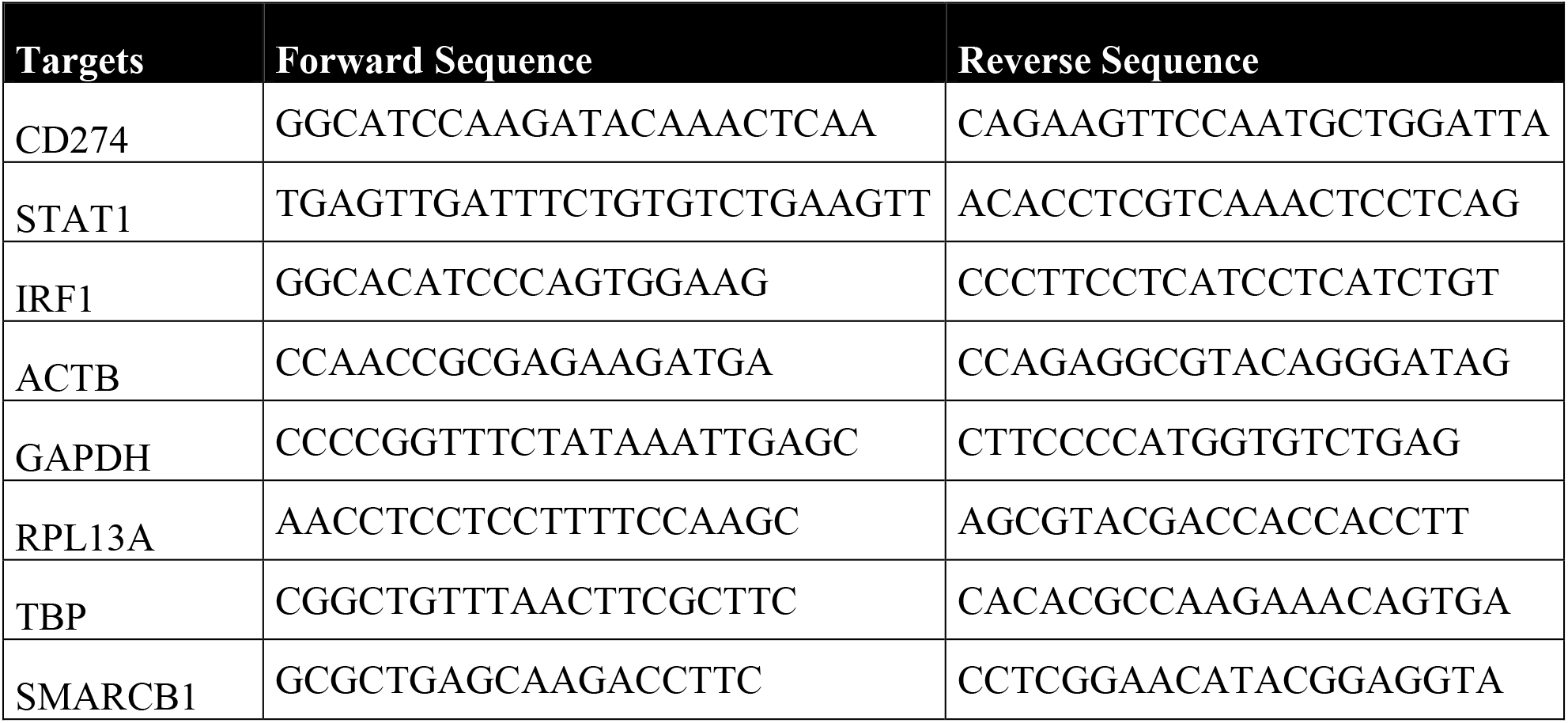
List of qPCR primers.

### Public data correlation analysis using TGCA and CCLE database

Spearman correlation for all selected genes were retrieved from co-expression studies using the Cancer Cell Line Encyclopaedia (Broad, 2019) and the TCGA chromophobe renal cell carcinoma (KICH) databases. All transcription factors were extracted using the “Full Human TFs” list from (Lambert et al., 2018). Scatter plots were made using Prism5. For the correlation with TFCP2L1, the epithelial and mesenchymal genes were retrieved from (Watanabe et al., 2019).

### Bulk RNA sequencing

RMC cell lines were analysed by RNA-seq under the different indicated conditions. After sequencing raw reads were pre-processed in order to remove adapter and low-quality sequences (Phred quality score below 20) using cutadapt version 1.10. and reads shorter than 40 bases were discarded. Reads were maping to rRNA sequences using bowtie version 2.2.8, were also removed. Reads were mapped onto the hg19 assembly of Homo sapiens genome using STAR version 2.5.3a. Gene expression quantification was performed from uniquely aligned reads using htseq-count version 0.6.1p1, with annotations from Ensembl version 75 and “union” mode. Only non-ambiguously assigned reads were retained for further analyses. Read counts were normalized across samples with the median-of-ratios method. Comparisons of interest were performed using the Wald test for differential expression and implemented in the Bioconductor package DESeq2 version 1.16.1. Genes with high Cook’s distance were filtered out and independent filtering based on the mean of normalized counts was performed. P-values were adjusted for multiple testing using the Benjamini and Hochberg method. Deregulated genes were defined as genes with log2(foldchange) >1 or <-1 and adjusted p-value <0.05.

### Analysis of bulk RNA-seq of patient samples

For RMC cohorts, raw counts were retrieved in excel format and normalized first by sequencing depth using DESeq2 sizefactors and then divided by median of gene length. Samples were clustered using the hclust function with “ward.D2” linkage function and visualized as heatmaps using pheatmap package v1.0.12. The deconvolution of immune and stromal cells was done using MCP-counter v1.2.0 (Becht et al., 2016). Sample compositions were also estimated by deconvolution from our single-cell data using the CIBERSORTx algorithm (Newman et al., 2019). Volcano plots were generated with ggplot2 v3.3.2. Gene set enrichment analyses were done with the GSEA software v3.0 using the hallmark gene sets of Molecular Signature Database v6.2. Gene Ontology analysis was done using DAVID (http://david.abcc.ncifcrf.gov/). Gene list intersections and Venn diagrams were performed by Venny.

### Protein extraction and Western blotting

Whole cell extracts were prepared by the standard freeze-thaw technique using LSDB 500 buffer (500 mM KCl, 25 mM Tris at pH 7.9, 10% glycerol (v/v), 0.05% NP-40 (v/v), 16mM DTT, and protease inhibitor cocktail). Cell lysates were subjected to SDS–polyacrylamide gel electrophoresis (SDS-PAGE) and proteins were transferred onto a nitrocellulose membrane. Membranes were incubated with primary antibodies in 5% dry fat milk and 0.01% Tween-20 overnight at 4 °C. The membrane was then incubated with HRP-conjugated secondary antibody (Jackson ImmunoResearch) for 1h at room temperature, and visualized using the ECL detection system (GE Healthcare). The references of all antibodies are available in Table 1.

### Chromatin immunoprecipitation and sequencing (ChIP-seq)

BRG1 ChIP experiments were performed on native MNase-digested chromatin. Between 10 to 20 × 10^8^ freshly harvested RMC2C cells bearing either SMARCB1 or mCHERRY and treated 2uM doxycycline for 48hrs were resuspended in 1.5 ml ice-cold hypotonic buffer (0.3M Sucrose, 60 mM KCl, 15 mM NaCl, 5 mM MgCl2, 0.1 mM EDTA, 15 mM Tris–HCl pH 7.5, 0.5 mM DTT, 0.1 mM PMSF, PIC) and cytoplasmic fraction was released by incubation with 1.5 ml of lysis-buffer (0.3M sucrose, 60 mM KCl, 15 mM NaCl, 5 mM MgCl2, 0.1 mM EDTA, 15 mM Tris–HCl pH 7.5, 0.5 mM DTT, 0.1 mM PMSF, PIC, 0.5% (vol/vol) IGEPAL CA-630) for 10 min on ice. The suspension was layered onto a sucrose cushion (1.2 M sucrose, 60 mM KCl, 15 mM NaCl, 5 mM MgCl2, 0.1 mM EDTA, 15 mM Tris–HCl [pH 7.5], 0.5 mM DTT, 0.1 mM PMSF, PIC) and centrifuged for 30 min 4°C at 4700 rpm in a swing rotor. The nuclear pellet was resuspended in digestion buffer (0.32Msucrose, 50 mM Tris–HCl [pH 7.5], 4 mM MgCl2, 1 mM CaCl2, 0.1 mM PMSF) and incubated with 10ul of Micrococcal Nuclease (NEB) for 7 min at 37°C. The reaction was stopped by addition of 20ul of EDTA 0,5M and suspension chilled on ice for 10 min. The suspension was cleared by centrifugation at 10,000 rpm (4°C) for 10 min and supernatant (chromatin) was used for further purposes. Chromatin was digested to around 80% of mono-nucleosomes as judged by extraction of the DNA and agarose gel electrophoresis. H3K27ac and MYC ChIP experiments were performed on 0.4% PFA-fixed chromatin isolated from RMC2C cells bearing either SMARCB1 or mCHERRY and treated 2uM doxycycline for 48hrs according to standard protocols as previously described (Strub et al., 2011). ChIP-seq libraries were prepared using MicroPlex Library Preparation kit v2 and sequenced on the Illumina Hi-seq 4000 as single-end 50-base reads (Laurette et al., 2020). Sequenced reads were mapped to the Homo sapiens genome assembly hg19 using Bowtie with the following arguments: -m 1 --strata --best -y -S -l 40 -p 2.

### ChIP-seq analysis

After sequencing, peak detection was performed using the MACS software (Zhang et al., 2008). Peaks were annotated with Homer using the GTF from ENSEMBL v75. Global clustering analysis and quantitative comparisons were performed using seqMINER (Ye et al., 2011). Super-enhancers were called with the python package Ranking Of Super Enhancers (ROSE) by ranking H3K27ac peaks either by H3K27ac, BRG1 or MYC signal.

De novo motif discovery on FASTA sequences corresponding to windowed peaks was performed using MEME suite (meme-suite.org). Motif correlation matrix was calculated with in-house algorithms using JASPAR database as described in (Joshi et al., 2017). Motif discovery from ChIP-seq peaks was performed using the RSAT peak-motifs algorithm (http://rsat.sb-roscoff.fr/peak-motifs_form.cgi).

Motif analysis Searching of known TF motifs from the Jaspar 2014 motif database at BRG1-bound sites was made using FIMO (Grant et al., 2011) within regions of 200 bp around peak summits, FIMO results were then processed by a custom Perl script which computed the frequency of occurrence of each motif. To assess the enrichment of motifs within the regions of interest, the same analysis was done 100 times on randomly selected regions of the same length as the BRG1 bound regions and the results used to compute an expected distribution of motif occurrence. The significance of the motif occurrence at the BRG1-occupied regions was estimated through the computation of a Z-score (z) with z = (x − μ)/σ, where: − x is the observed value (number of motif occurrence), − μ is the mean of the number of occurrences (computed on randomly selected data), − σ is the standard deviation of the number of occurrences of motifs (computed on randomly selected data). The source code is accessible at https://github.com/slegras/motif-search-significance.git.

### Statistics

All experiments were performed in biological triplicates, unless stated otherwise in the figure legends. All tests used for statistical significance were calculated using Prism5 and are directly indicated in the figure legends (****: p < 0.0001, ***: p < 0.001, **: p < 0.01, *: p < 0.05, ns: p > 0,05).

### Data availability

Source data for this paper are available from the authors upon reasonable request. All sequencing data reported here have been submitted to the GEO database under accession number GSE181001.

## Acknowledgements

We thank all the staff of the IGBMC common facilities in particular Betty Heller and Patricia Wagner from Cell Culture, Claudine Ebel and Muriel Philipps from Flow Cytometry and Dr. Paola Rossolillo and Karim Essabri of the molecular biology facility. We also would like to thank the Institut Curie facilities and in particular Dr Pascale Philippe-Chomette, Pr Michel Peuchmaur, Dr Yves Allory and Dr Pascale Maille for proving the primary specimen for the PDX experiments.

This work was supported by institutional grants from the Centre National de la Recherche Scientifique, the Institut National de la Santé et de la Recherche Médicale, the Université de Strasbourg, the Association pour la Recherche contre le Cancer (CR, contract number PJA 20181208268), the Ligue Nationale contre le Cancer, the Institut National du Cancer, the ANR-10-LABX-0030-INRT French state fund through the Agence Nationale de la Recherche under the frame programme Investissements d’Avenir labelled ANR-10-IDEX-0002-02.

The IGBMC high throughput sequencing facility is a member of the “France Génomique” consortium (ANR10-INBS-09-08). FB was supported by grants from the INCA for the scRNAseq program (INCA PRTK19-2020-036) and the Saint Bladrick Foundation. ID is an ‘équipe labellisée’ of the Ligue Nationale contre le Cancer. BHV was supported by fellowships from the ANR, the Ligue Nationale contre le Cancer.

Conceptualization, BHV, ID, GGM; Methodology, BHV, GD, AH, MR, ARH, JG, JT, PB, PM, XS, DS, RB, JEK, FB, NMT, ID, GGM; Formal Analysis, BHV, GD, ID, GGM. Investigation, BHV, GD, AH, MR, ARH, JG, JT, PB, PM, XS, HL, TT, VL, DS, RB, JEK, FB, NMT, ID, GGM. Bioinformatics: GD, BHV, XS; Writing – Original Draft, BHV, ID, GGM; Writing – Review and Editing, BHV, GD, ID, GGM. Investigation, BHV, GD, AH, MR, ARH, JG, JT, PB, PM, XS, HL, TT, VL, DS, RB, JEK, FB, NMT, ID, GGM; Resources, PM, XS, HL, TT, VL, DS, RB, FB, NMT, ID and GGM; Supervision: ID, GGM

## Legends to Figures

**Supplementary Figure 1.**
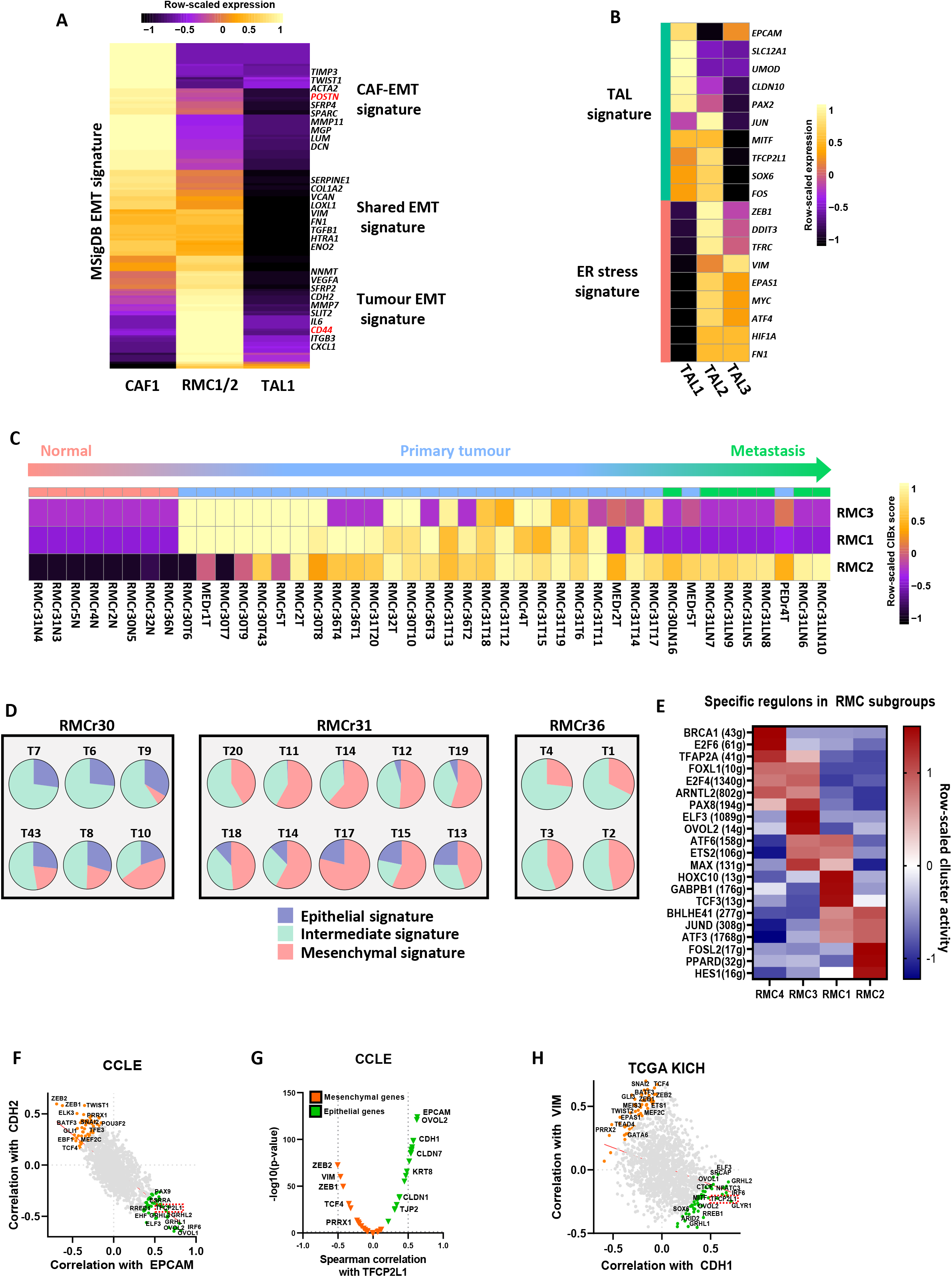
**A.** Pseudo-bulk heatmap showing expression of the MSigDB Hallmark EMT signature (183 genes) in RMC1/2 and CAF1 cells from the treated tumour. **B.** Pseudo-bulk heatmap showing heterogeneous expression of selected TAL identity markers, mesenchymal and ER stress genes in all TAL clusters. **C.** Deconvolution of RMC specific signatures as calculated by CIBERSORTx on bulk RNA-seq from sections of RMC primary tumours, lymph node metastasis and normal adjacent tissues. **D.** Pie charts representing intratumoural heterogeneity of RMC signatures using multi-region RNA sequencing of primary RMC tumours (n=3). Note that relative proportions (in %) were inferred by CIBERSORTx using our scRNA-seq normalized merge**. E.** SCENIC analysis of normalized merge of treated and naive RMC samples revealing specific regulons of all RMC clusters. **F-G.** Pearson correlation analysis of 1683 human transcription factors with selected genes in CCLE database (F) and in TGCA chromophobe RCC samples (G). Note the positive correlation of TFCP2L1 expression level with epithelial markers along with other Grainyhead family members, OVOL1/2 and MITF. **H.** Pearson correlation analysis of TFCP2L1 expression with a set of epithelial and mesenchymal genes in CCLE database.

**Supplementary Figure 2.**
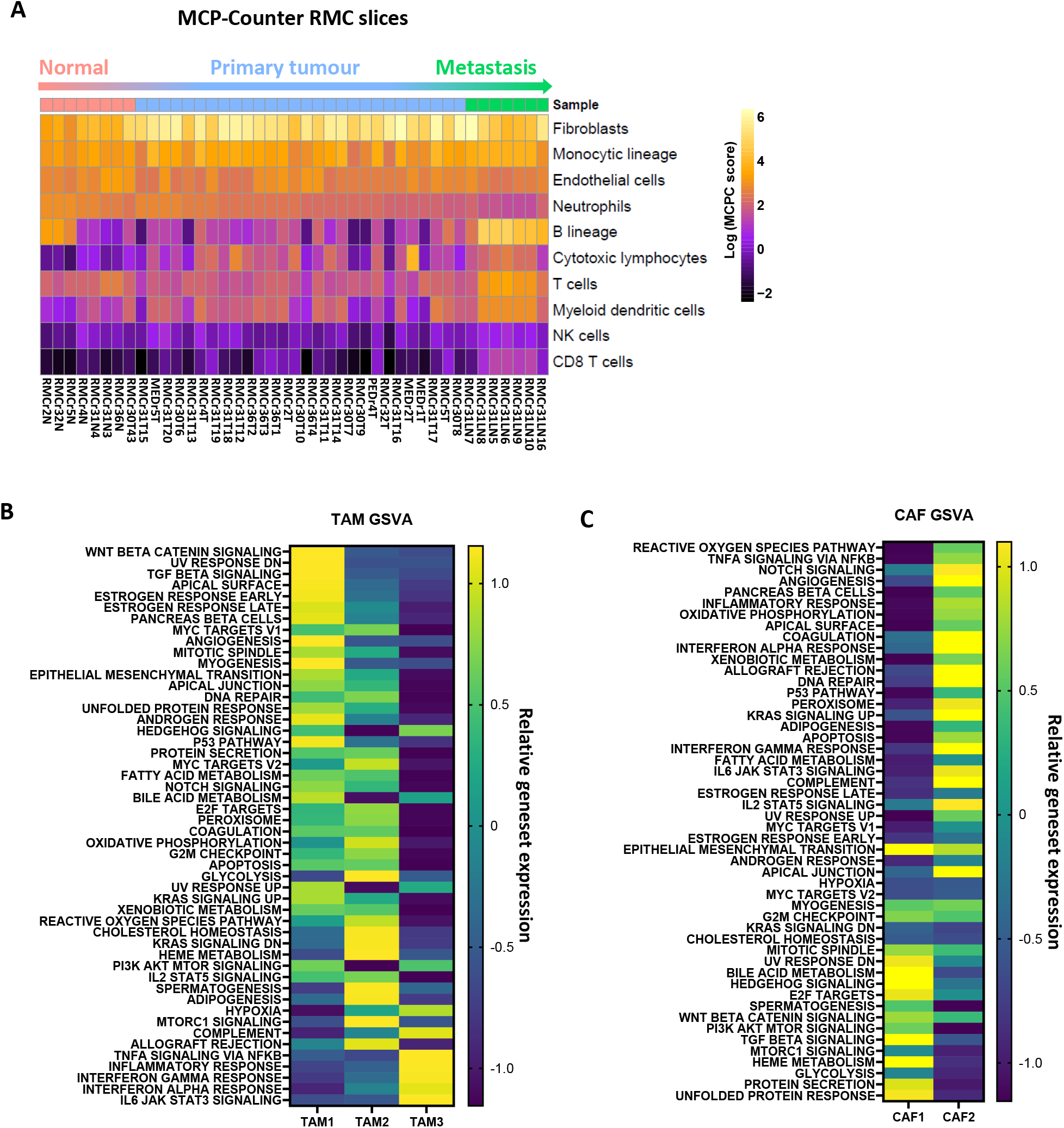
**A.** Heatmap revealing relative abundance of immune and stromal cell infiltrates scores in sections of the RMC cohort as inferred by MCP-Counter. Note the strong enrichment scores of fibroblasts and monocytic lineage cells relative to other cell populations. **B.** GSVA showing enrichment of Hallmarks genesets according to the TAM subclusters. **C.** GSVA showing enrichment of selected Hallmarks genesets in the CAF subclusters.

**Supplementary Figure 3.**
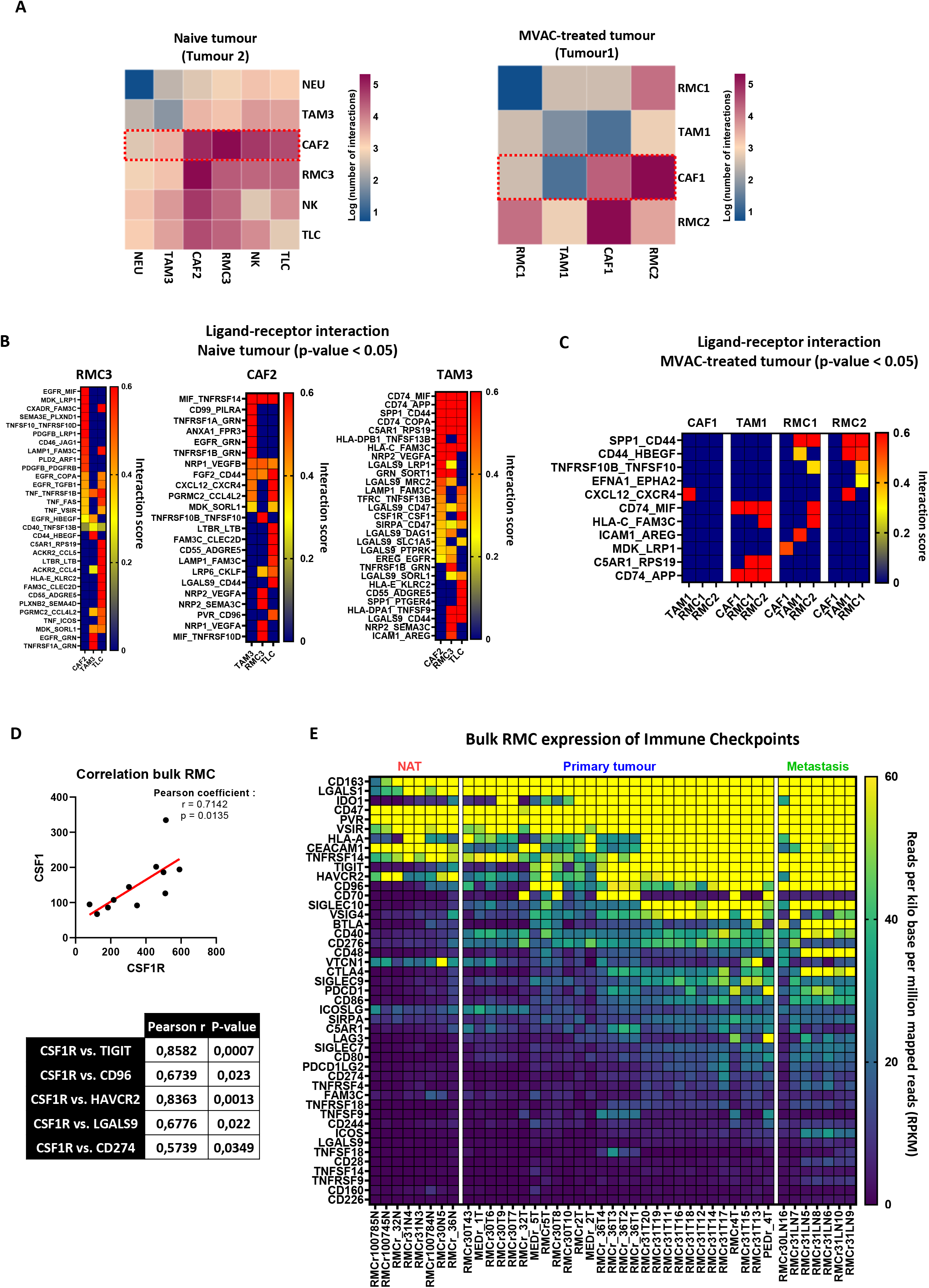
**A.** Heatmap showing ligand-receptor pair interactions between cells from the treated and the naive RMC samples as inferred by CellPhoneDB. **B.** Heatmap showing major cross-talks in the naive RMC sample between RMC3, CAF2, TAM3 and TLC cell populations. Note that pairs with p < 0.05 are shown on heatmaps using the Ligand-receptor (LR) interaction score provided by the program as ranking. **C.** Heatmap of major cross-talks (ranked by LR interaction score) in the treated RMC sample between CAF1, TAM1, RMC1 and RMC2 cell populations. Note that only significant pairs are shown. **D.** Pearson correlation r coefficient for expression level of CSF1R, its ligand CSF1 and other markers of immunosuppresion in bulk RMC primary tumours (MDACC, n=11). **E.** RPKM expression values of known immune checkpoint proteins using all RNAseq of NAT, RMC primary tumours and lymph node metastasis (n=44).

**Supplementary Figure 4.**
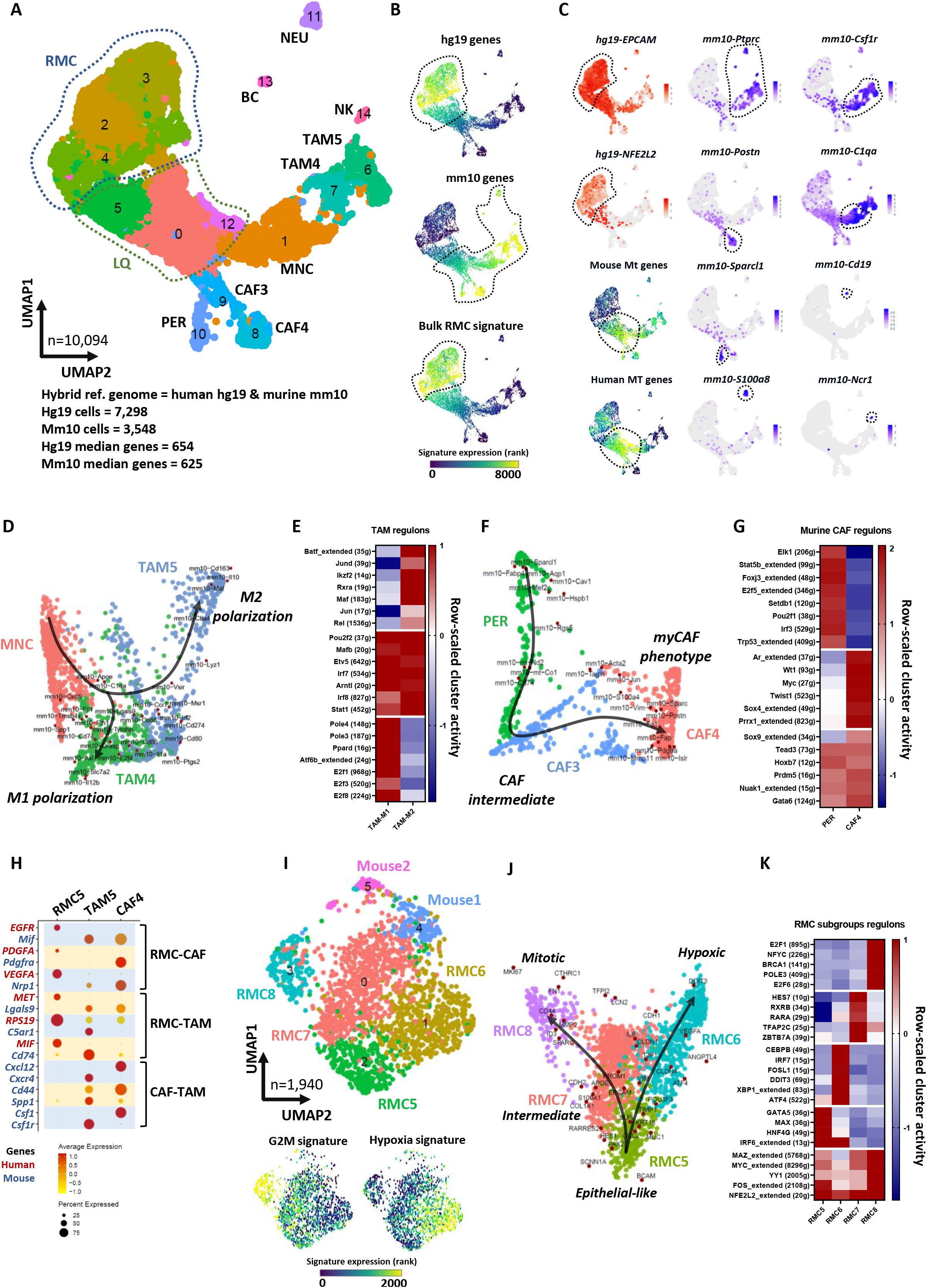
**A.** UMAP plot of scRNA-seq from the RMC PDX (IC-pPDX-132) representing cell clusters as calculated by Seurat at a resolution of 0.3. Clusters were identified using hallmark genes shown in Fig. 3B/C. *Abbreviations:* RMC : Renal medullary carcinoma cells; LQ : low quality cells; PER : pericytes; CAF3/4 : cancer-associated fibroblasts; MNC : monocytes; TAM4/5 : tumour-associated macrophages; NK: natural killers; BC: B-cells; NEU : neutrophils. **B.** UMAP projection of selected gene signatures. Human and murine signatures were established using differential gene nomenclature. **C.** UMAP projection of marker genes and mitochondrial gene signatures. Human and murine mitochondrial gene signatures were established using differential gene nomenclature. **D.** SWNE trajectory analysis of murine TAM clusters and their putative monocyte cell-of-origin showing the differential polarization of TAM cell populations. **E.** SCENIC analysis of murine TAM subclusters**. F.** SWNE trajectory analysis of murine CAF clusters and their putative pericyte cell-of-origin, showing the differential polarization of TAM cell populations. **G.** SCENIC analysis of murine pericyte and CAF clusters. **H**. Dot plot showing average expression of selected RMC ligand-receptor pairs identified previously in Figure 3I. Note that human genes are coloured in red, and murine genes in blue. **I**. UMAP representing PDX RMC subclusters as identified by Seurat using a resolution of 1 (upper panel), Average expression of selected MSigDB gene signatures (lower panel). **J.** SWNE trajectory analysis of RMC cells using markers of each cluster. **K.** SCENIC analysis of RMC subclusters.

**Supplementary Figure 5.**
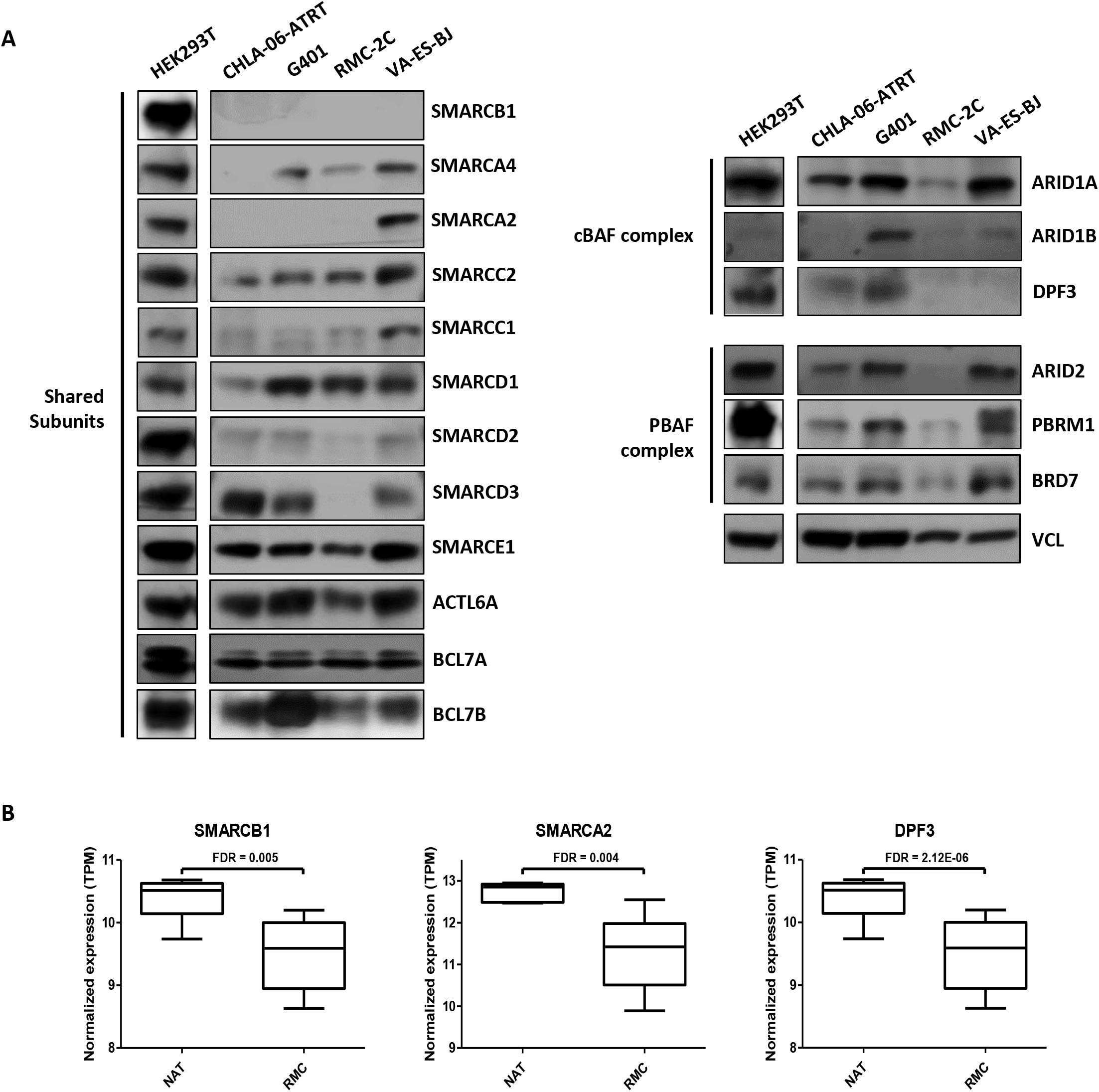
**A.** Immunoblots revealing expression of SWI/SNF subunits in RMC2C cells, 3 additional SMARCB1-deficient lines and HEK293T cells. HEK293T: immortalized human embryonic kidney cells; CHLA-06-ATRT: atypical teratoid/rhabdoid tumour cell line; G401: malignant rhabdoid tumour cell line; VA-ES-BJ: epithelioid sarcoma cell line. Loading normalisation: VCL. **B.** Box plots showing the expression of a selection of SWI/SNF genes in RMC and NAT, with associated FDR value for statistical significance.

**Supplementary Figure 6.**
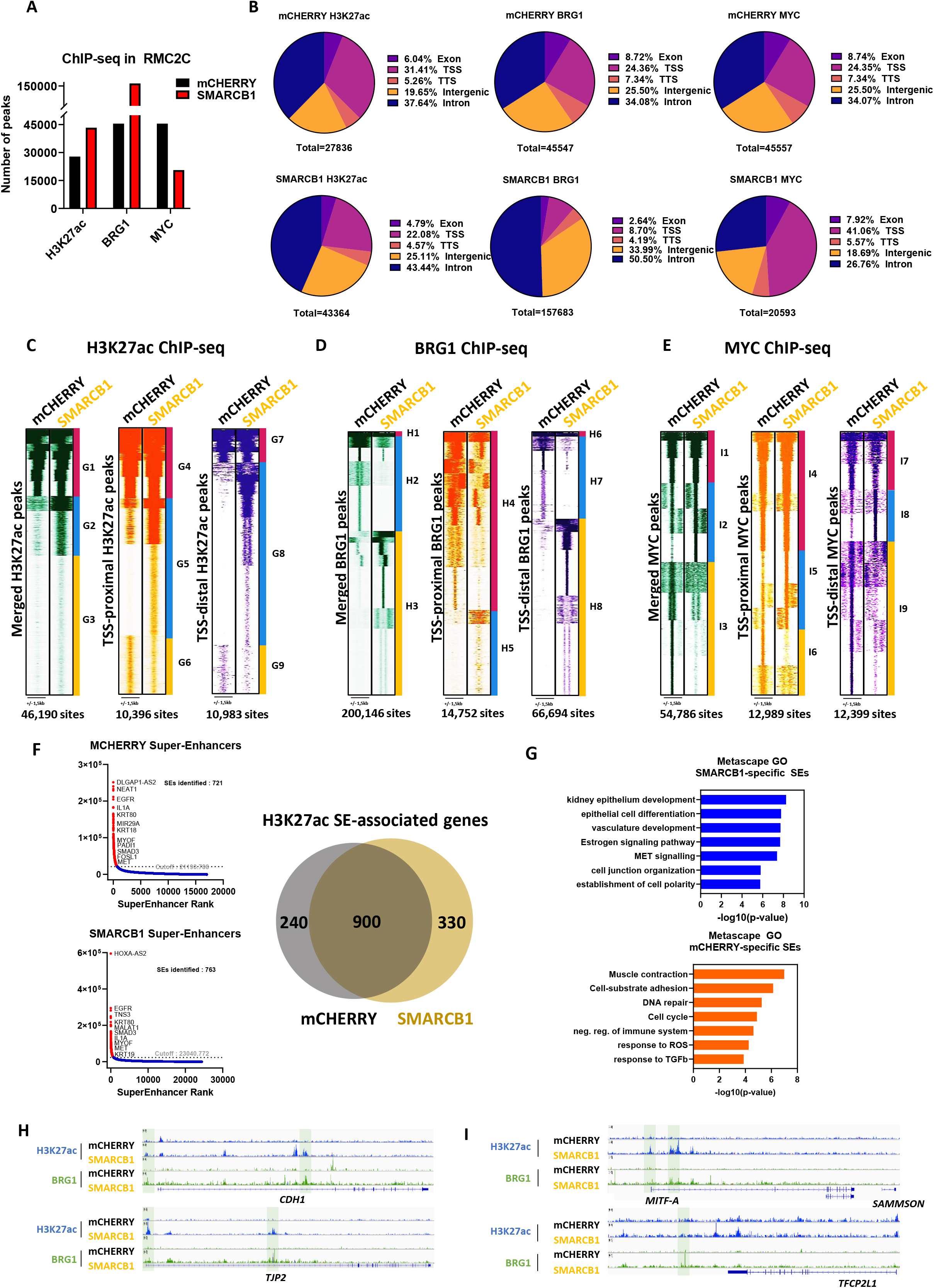
**A.** Number of peaks of H3K27ac, BRG1 and MYC in SMARCB1 or mCHERRY-expressing cells as quantified by the MACS algorithm. **B.** Pie charts showing the relative distribution of H3K27ac, BRG1 and MYC peaks on defined genome elements. **C-E**. Read density maps of H3K27ac (C), BRG1 (D), MYC (E) peaks in SMARCB1- or mCHERRY-expressing cells using either all merged, TSS-proximal or TSS-distal sites as a reference. **F.** ROSE identification of H3K27ac Super-Enhancers (SE) in RMC2C cells expressing SMARCB1 or mCHERRY (left), and Venn diagram of shared and specific SE-associated genes (right). **G.** Ontology enrichment analysis of mCHERRY-and SMARCB1-specific SE-associated genes. **H-I**. UCSC genome track snapshots showing BRG1 and H3K27ac at proximal and distal regulatory elements of selected relevant genes.

**Supplementary Figure 7.**
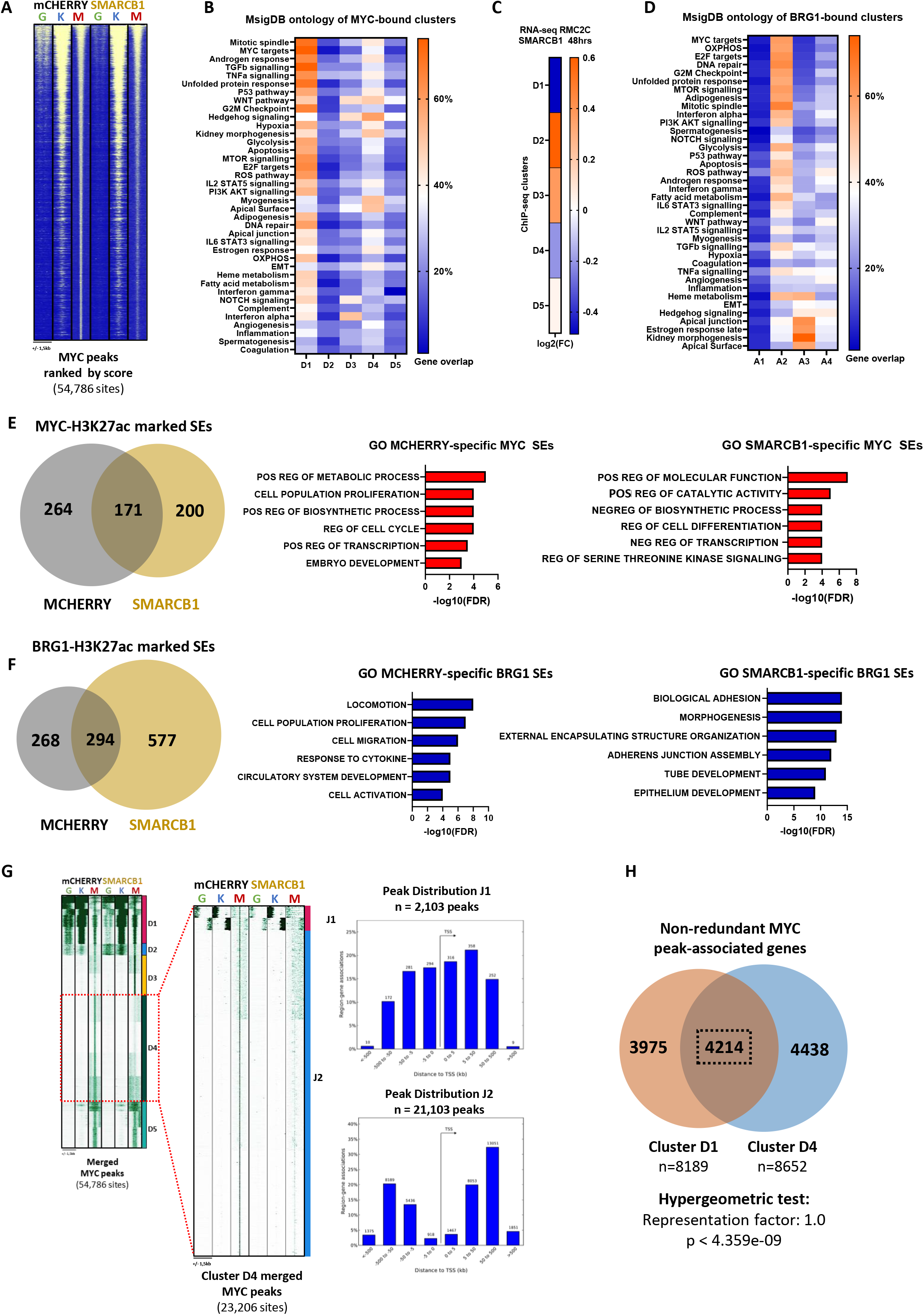
**A.** Tornado read density maps showing BRG1, H3K27ac and MYC sites ranked decreasingly by MYC peak score. **B.** Percentage of genes associated with MYC clusters as defined by HOMER in the indicated GSEA Hallmark Genesets. **C.** Changes in expression of genes associated with each MYC cluster upon SMARCB1 expression. **D.** Percentage of genes associated with BRG1 clusters as defined by HOMER in the indicated GSEA Hallmark Genesets. **E-F.** Venn diagram of SE-associated genes defined by MYC peak score (D) or BRG1 peak score (E) revealing common and specific SEs in SMARCB1- and mCHERRY-expressing RMC2C cells (left) and associated ontology analysis of SMARCB1- and mCHERRY-specific SE-associated genes (right). **G.** Read density maps showing sub-clustering of MYC D4 sites, with their peak distribution as calculated by GREAT (middle), and the associated ontology analysis of associated genes. **H.** Venn diagram of non-redundant MYC-bound genes found in clusters D1 and D4.

**Supplementary Dataset 1.** Markers defining cell clusters of the treated tumour, naive tumour and PDX (IC-pPDX-132) cells as calculated by Seurat FindMarkers algorithm.

**Supplementary Dataset 2.** Bulk RNA-seq of NAT, RMC primary tumours and lymph node metastases (n=44) from the MDACC cohort.

**Supplementary Dataset 3.** RNA-seq of RMC2C and RMC219 cells at 12hrs and 48hrs after SMARCB1 re-expression.

## Notes

The authors declare no potential conflicts of interest

### Competing Interest Statement

The authors have declared no competing interest.

